# Inferring single-trial neural population dynamics using sequential auto-encoders

**DOI:** 10.1101/152884

**Authors:** Chethan Pandarinath, Daniel J. O’Shea, Jasmine Collins, Rafal Jozefowicz, Sergey D. Stavisky, Jonathan C. Kao, Eric M. Trautmann, Matthew T. Kaufman, Stephen I. Ryu, Leigh R. Hochberg, Jaimie M. Henderson, Krishna V. Shenoy, L. F. Abbott, David Sussillo

## Abstract

Neuroscience is experiencing a data revolution in which simultaneous recording of many hundreds or thousands of neurons is revealing structure in population activity that is not apparent from single-neuron responses. This structure is typically extracted from trial-averaged data. Single-trial analyses are challenging due to incomplete sampling of the neural population, trial-to-trial variability, and fluctuations in action potential timing. Here we introduce Latent Factor Analysis via Dynamical Systems (LFADS), a deep learning method to infer latent dynamics from single-trial neural spiking data. LFADS uses a nonlinear dynamical system (a recurrent neural network) to infer the dynamics underlying observed population activity and to extract ‘de-noised’ single-trial firing rates from neural spiking data. We apply LFADS to a variety of monkey and human motor cortical datasets, demonstrating its ability to predict observed behavioral variables with unprecedented accuracy, extract precise estimates of neural dynamics on single trials, infer perturbations to those dynamics that correlate with behavioral choices, and combine data from non-overlapping recording sessions (spanning months) to improve inference of underlying dynamics. In summary, LFADS leverages all observations of a neural population’s activity to accurately model its dynamics on single trials, opening the door to a detailed understanding of the role of dynamics in performing computation and ultimately driving behavior.

---

Increasing evidence suggests that in many brain areas, such as the motor and prefrontal cortices, the activity of large populations of neurons, termed the neural population state, is often well-described by low-dimensional dynamics [e.g. (Afshar et al. 2011; Harvey, Coen, and Tank 2012; Churchland et al. 2012; Mante et al. 2013; Kaufman et al. 2014; Sadtler et al. 2014; Pandarinath et al. 2015; Carnevale et al. 2015; Kobak et al. 2016a)]. Recovering these dynamics on single trials is essential for illuminating the relationship between neural population activity and behavior, and for advancing therapeutic neurotechnologies such as closed-loop deep brain stimulation and brain-machine interfaces. However, recovering population dynamics on single trials is difficult due to trial-to-trial variability (e.g. in behavior or arousal) and fluctuations in the spiking of individual neurons. Even with dramatic increases in the numbers of neurons that can be simultaneously recorded using multichannel electrode arrays or optical imaging, accurately recovering population dynamics from single trials remains a significant challenge for data-analysis methods.

Standard analyses sacrifice single-trial information for the sake of better estimates of trial-averaged neural states (Ahrens et al. 2012; Churchland et al. 2012; Mante et al. 2013; Kobak et al. 2016b). Techniques for extracting neural population states from single trials exist and are in use, but they typically make simplifying assumptions by modeling the underlying population dynamics as having independent underlying factors (Yu et al. 2009), as being linear (Macke et al. 2011; Kao et al. 2015; Aghagolzadeh and Truccolo 2014; Zhao and Park 2017; Y. Gao et al. 2016) or as being switched linear (Petreska et al. 2011; Linderman et al. 2017). Here we introduce a novel machine learning method based on nonlinear artificial recurrent neural networks (RNNs), termed Latent Factor Analysis via Dynamical Systems (LFADS, “*ell-fads*”). LFADS is fully nonlinear, provides estimates of inputs from areas not being recorded, and can be applied to extract shared structure from data collected from different populations across multiple recording sessions.

LFADS is based on the assumption that spiking activity on a single trial of a task depends on: 1) underlying dynamics characteristic of the brain area(s) being recorded; 2) trial-specific initial conditions for those dynamics that reflect both external and internal states of the subject; 3) effects of unmeasured inputs from other brain areas, including those arising from unexpected changes in task structure or internal state during a trial, and 4) Poisson spiking variability. In LFADS (**Fig. 1a**), the underlying dynamics (assumption 1) are generated by an RNN (the “generator” network). Dynamic “factors” are extracted from this RNN and are used to generate (and thereby infer) firing rates of the recorded neurons. The inferred firing rates generate action potentials through a Poisson process (assumption 4). Initial conditions and input for the generator network (assumptions 2 and 3) are extracted from spiking data for each trial by additional RNNs (the “encoder” and “controller” networks). Yet, beyond binned spike sequences, no other trial-specific information is supplied to the model.

**Figure 1.**
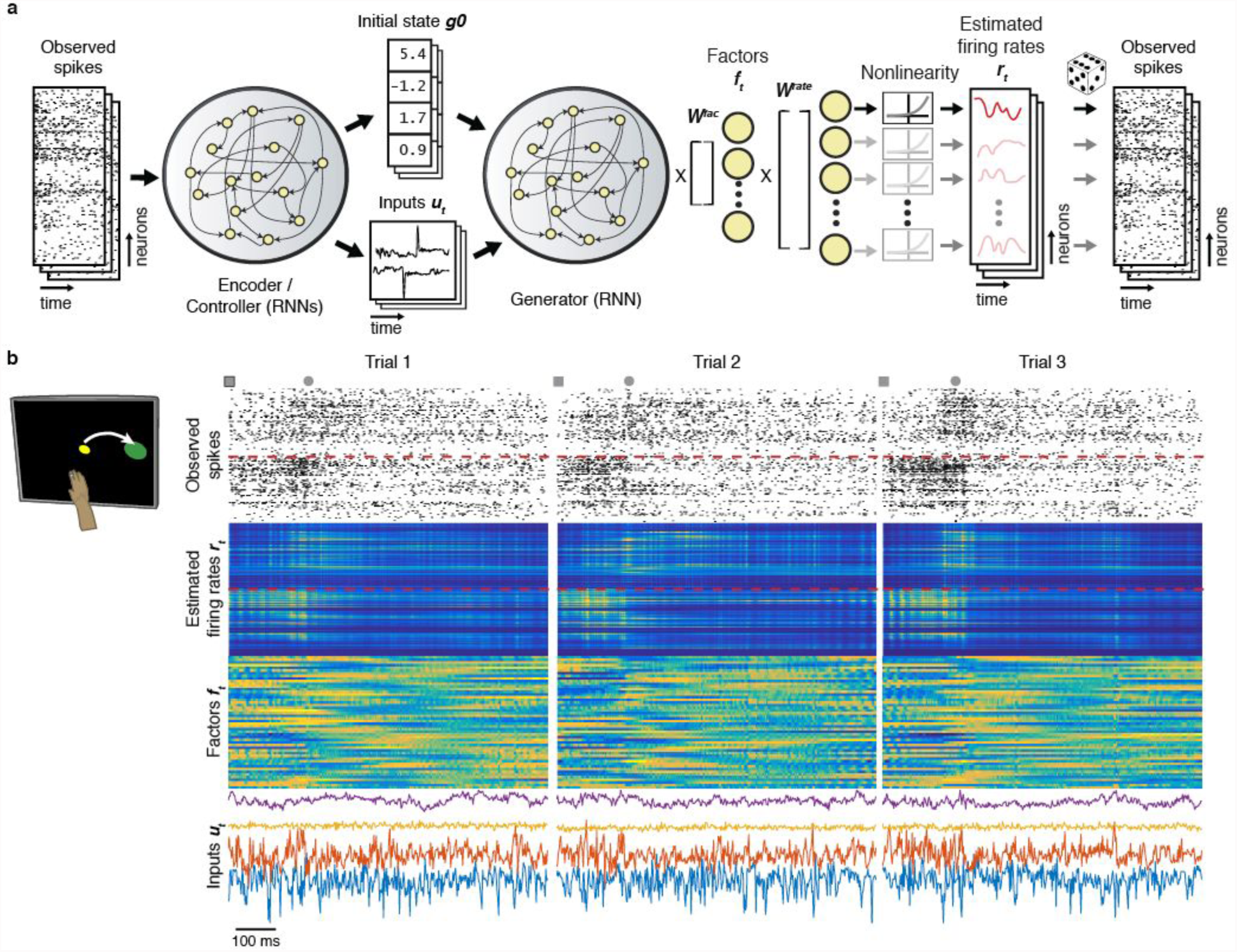
LFADS is a generative model that assumes that observed single-trial spiking activity is generated by an underlying dynamical system. (**a**) LFADS takes a given recording (far left), reduces is to a latent code consisting of an inferred initial condition and inputs (middle), and then attempts to estimate firing rates that are consistent with the observed data (right) from that latent code. I.e. LFADS auto-encodes the trial via a sequential auto-encoder. Working from right to left in the panel, for the *i^th^* neuron, LFADS estimates firing rates at time *t*, *r_t,i_*, for each of 192 channels, and the observed spike counts are assumed to be Poisson distributed count observations of these underlying firing rates. The firing rates are linear readouts from a set of low-dimensional factors ***f_t_*** (50) via a readout matrix ***W^rate^***. The factors are defined as linear readouts from a dynamical generator (an RNN), via a readout matrix ***W^fac^***. Activity of the generator is determined by its per-trial components consisting of an initial state (***g_0_***) and 4 time-varying inferred inputs (***u_t_***), and its recurrent connectivity, which is fixed for all trials. Both ***g_0_*** and ***u_t_*** are determined from a single-trial recording via a set of encoder and controller RNNs. (**b**) Example spiking activity recorded from M1 and PMd (red dashed line separates M1 above from PMd below) as a monkey performed a reaching task, as well as the corresponding firing rates ***r_t_***, factors ***f_t_***, and inferred inputs ***u_t_***, all inferred by LFADS (3 example trials are shown). Squares denote time of target onset, and circles denote time of movement onset.

The strength of this approach lies in exploiting the capacity of nonlinear RNNs to reproduce complex temporal patterns of activity that underlie the neural data. LFADS combines information obtained from all the recorded neurons on all trials, as well as single-trial spiking data, to produce de-noised firing rates for each neuron on each trial. This contrasts with conventional binning or filtering of spike trains, which computes rates solely from the activity on the single trial being analyzed. LFADS is forced to find low-dimensional dynamics that explain the recorded data because the number of dynamic factors in the model is constrained. This is consistent with repeated empirical observations that the dimensionality of neural population activity in areas like motor and prefrontal cortices is, in several cases, much lower than the number of recorded neurons [(Churchland et al. 2012; Mante et al. 2013; Kato et al. 2015; Kaufman et al. 2016); see (P. Gao and Ganguli 2015) for a full discussion].

Here we apply LFADS to a variety of datasets from rhesus macaque motor and pre-motor cortices using both chronic electrode arrays and acute linear array recordings as well as human motor cortex using chronic electrode arrays (datasets and recording methodologies are outlined in **Online Methods Table 2**; macaque data were previously recorded at Stanford University). We show that firing rates extracted by LFADS can be used to estimate behavioral variables (e.g. reaching kinematics) significantly more accurately than other techniques, both standard and state-of-the-art. We also show that the dynamics inferred by LFADS capture features of the data on multiple timescales, including those related to behavior and faster timescales associated with local field potential oscillations. Further, we demonstrate that LFADS can combine data from non-overlapping recording sessions, sampling from separate neural populations, to improve its performance on each session. Finally, we demonstrate the ability of LFADS to infer inputs to a neural circuit by analyzing data from an arm-reaching task involving a mid-trial perturbation.

## Results

### Overview of LFADS

LFADS is a sequential adaptation of a variational auto-encoder (Kingma and Welling 2013) constructed by maximizing a lower bound on the the likelihood of the observed spiking activity being produced by the generator network, across all trials except those held out for cross-validation purposes. Penalties are imposed on the complexity of the initial conditions and inferred inputs to ensure that the generator model explains as much of that data as possible [see e.g. (Gregor et al. 2015)]. Parameters are learned using standard deep learning methodologies, namely stochastic gradient descent and backpropagation (full details of the model and training procedure are given in Online Methods, and associated source code is available).

Working from output (right) to input (left) in (**Fig. 1a**), LFADS models a vector of single-trial spiking observations as stochastic (Poisson) spike counts generated from a vector of underlying firing rates ***r_t_***, at time *t*. For neuron *i*, the LFADS-inferred firing rate *r_t,i_* provides a de-noised rate for its observed spiking activity on a trial-by-trial basis. The firing rates are obtained by multiplying a vector of dynamic factors ***f_t_*** by a readout matrix ***W^rate^*** and then computing an exponential function of the resulting quantity. The vector of dynamic factors is determined by multiplying the vector of activities ***g_t_*** of the generator RNN by a matrix ***W^fac^***. The activities of the units of the generator RNN depend on three elements: a trial-specific initial state vector ***g_0_*** (one for each trial), a trial-specific vector of time-varying inputs ***u_t_***, and the parameters defining the connections of the network (which are fixed across trials after training). The inferred initial state ***g_0_*** and inputs ***u_t_*** are provided by linear readouts of the activities of additional RNNs, the encoder and controller. To compute these for a given trial, the encoder and controller RNNs receive input consisting of the vector of recorded spike counts within specified bins on that trial. To better model the trials, the LFADS encoders run through the trial both backwards and forwards to determine the values of ***g_0_*** and the inputs, meaning that when generating the trial at any time *t*, LFADS has access to data before and after *t*. As a consequence, the latent variables inferred by LFADS are more accurate, but this also means that the inputs inferred by LFADS are acausal with respect to the timing of external events.

In summary, once the model has been trained, recorded binned spike counts from a specific trial are fed into the LFADS encoder, which infers initial conditions and inputs for that trial **(Fig. 1a)**. The LFADS encoder effectively compresses the information contained in the spiking data for each trial into a “latent code”, the values of the initial conditions and inputs to the generator. From this compressed code, the generator RNN infers the dynamic factors and firing rates of all the recorded neurons across time for the encoded trial (in **Supp. Fig. 1** we apply LFADS to a 1-D pendulum to show, for a simple dynamical system, how LFADS operates). Thus, LFADS turns time series of single-trial recorded spike counts into inferred inputs, low-dimensional dynamic factors and the underlying firing rates which generated the observed spikes (see Online Methods for further details).

To illustrate the operation of LFADS, we begin by training LFADS models on array data (single-trial spiking activity) from macaque primary motor (M1) and premotor (PMd) cortices, recorded while a monkey made reaching movements in a target acquisition task (for details on training LFADS models, see Online methods, and **Online Methods Table 1**, for all model hyperparameters). The data sequences analyzed were 800 ms long and aligned to target onset at the start of each trial (i.e., the time when the target for the upcoming reach was presented). Inferred firing rates, dynamic factors, and inputs, for three example trials reveal several interesting features (**Fig. 1b**). The firing rates exhibit the expected relationship to the spiking activity (upper two panels in **Fig. 1b**), and they also display quite prominent oscillations that are most apparent in the rates (but not in the spikes) near the beginning of each trial (detailed in **Fig. 4**). In addition to this fast-timescale structure, further analysis in the next section reveals a second set of oscillations at a slower, behavioral timescale (detailed in **Fig. 3**). Thus, LFADS uncovered temporal dynamics on multiple timescales. Further, LFADS inferred separate firing rate dynamics for PMd and for M1 activity without being provided with information on the identity of any of the recorded channels (upper two panels in **Fig. 1b**, M1 above red dashed line, PMd below). The dynamic factors (third panel) compress the dynamics down to 50 dimensions. The 4-dimensional inferred inputs (bottom traces in **Fig. 1b**) affect the generator RNN, and also show variation on multiple timescales. In these examples the inferred input is being used to explain LFP oscillations (detailed in **Fig. 4**).

We have assessed the validity and accuracy of firing rates, dynamic factors and inputs, inferred by LFADS from simulated data for which the ground-truth is known, **Supp. Figs 2, 3, 6-8 and Supp. Table 1**. These studies show extensive comparisons of LFADS to a number of state-of-the-art machine-learning based techniques [GPFA (Yu et al. 2009); PfLDS (Y. Gao et al. 2016); and vLGP (Zhao and Park 2017)]. However, assessing the quality and validity of results on real data, like those shown in **Fig. 1b**, is difficult because the ground truth is either unknown (in the case of inferred inputs) or non-existent (because single-trial “instantaneous” firing rates are abstractions). As a first pass at addressing the validity of LFADS on real data, we can ask whether LFADS dynamic factors are predictive of held out data. Because the dynamic factors extracted by LFADS reflect the full neural population dynamics, they should be predictive for neurons that were not used to train the model (i.e., held-out neurons). Indeed, the low-dimensional representation produced by LFADS dynamic factors provides substantially improved single-trial likelihood of held-out neurons’ spikes over alternate techniques (**Supp. Fig. 4**). We exploit this feature when we “stitch” together data from different sessions (**Fig. 5**).

In the following sections, we test the validity of LFADS-inferred firing rates in a number of ways: we verify that they exhibit features seen in trial-averaged analysis (**Fig. 3**); predict details of behavior (**Figs. 2 & 5**); and correlate with local field potentials (LFPs), another population activity measure (**Fig. 4**). To establish the validity of inferred inputs, we show that they are informative about task perturbations and about the behavioral response they evoke (**Fig. 6**).

**Figure 2.**
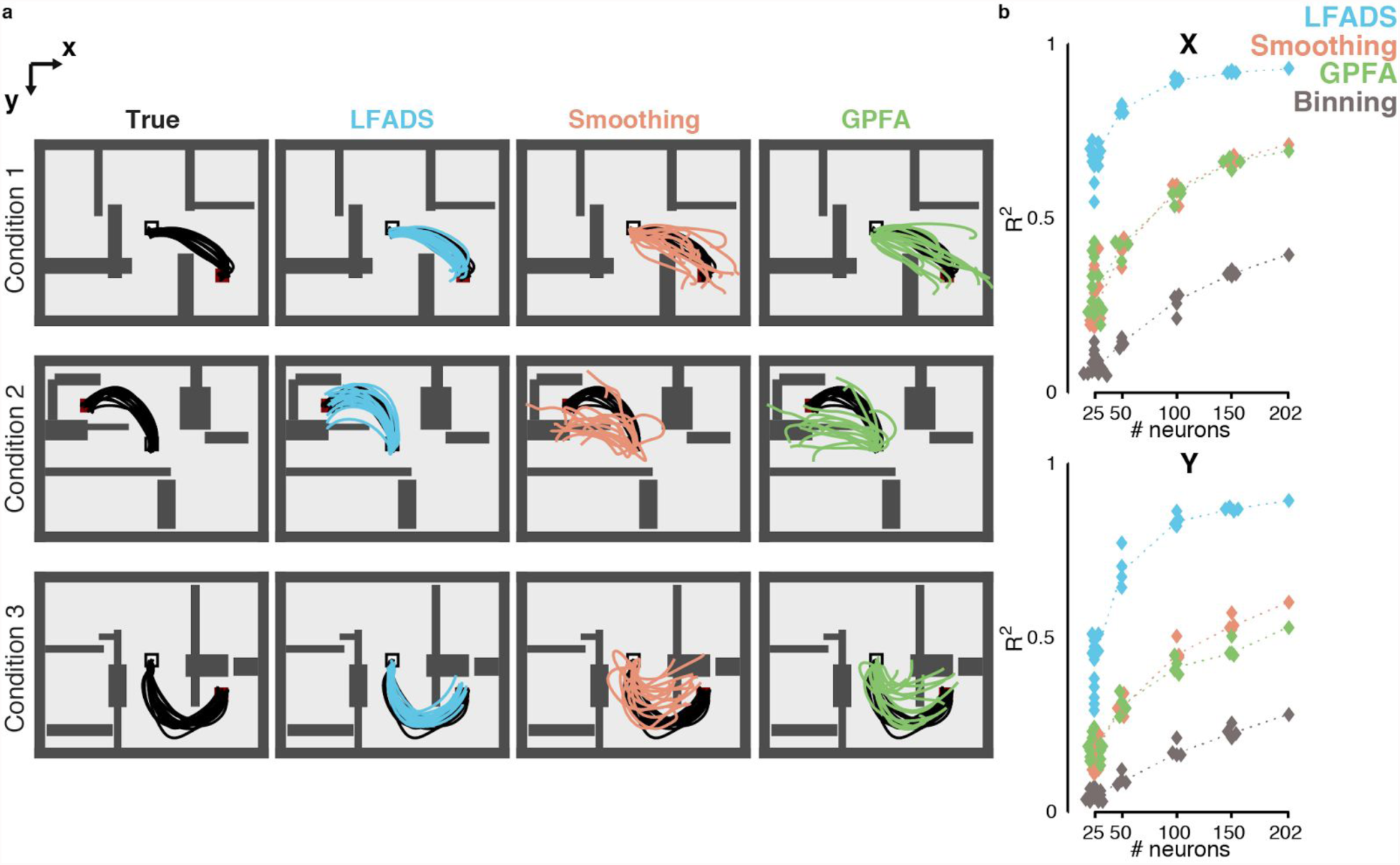
Application of LFADS to decoding reaching kinematics in a “Maze” reaching task. (**a**) A monkey was trained to perform arm reaching movements to guide a cursor in a 2-D plane from a starting location (center of the workspace) to peripheral targets. Virtual barriers in the workspace facilitated instruction of curved (or straight) reaches on a per-condition basis. Each row shows an example condition (3 shown, of 108 total). First column: true reach trajectories (black traces, 15 example trials per condition). Columns 2-4: examples of cross-validated reconstruction of these trajectories using Optimal Linear Estimation applied to the neural data, which was first de-noised either via LFADS, by smoothing with a Gaussian filter (40 ms s.d.), or using GPFA to reduce its dimensionality. (**b**) Decoding accuracy was quantified by measuring variance explained (R^2^) between the true and decoded velocities for individual trials across the entire dataset (2296 trials), for all three techniques and additionally for simple binning of the neural data. Accuracy was also measured for random sub-samples from the full neural population of 202 neurons (12 draws were tested for 25 neuron populations, and 4 draws each for the 50, 100, and 150 neuron populations). Dotted lines connect the median R^2^ values for each population size.

### Movement decoding and motor cortex dynamics

We applied LFADS to a population of 202 neurons (single units) simultaneously recorded from M1/PMd during a “Maze” task (see Online Methods) in which monkeys made a variety of straight and curved reaches through different arrangements of virtual barriers (**Fig. 2a**; dataset contained ~2300 individual reach trials). As in all of the examples we show, LFADS received no information about task instructions (the barrier patterns) or behavioral parameters. Estimates of hand velocities were computed from firing rates inferred by LFADS using cross-validated optimal linear estimation [OLE; (Salinas and Abbott 1994)]. We compared the accuracy of these velocities with results obtained by applying OLE to single-trial firing rates or low-dimensional state estimates inferred by other widely used and high-performing techniques. Using the full population of 202 neurons, decoding using LFADS estimated rates dramatically outperformed results obtained by binning or filtering spike trains, or by using Gaussian Process Factor Analysis [GPFA; (Yu et al. 2009)] (**Fig. 2a,b**; average R^2^ of 0.91 across the dataset, vs. 0.66, 0.61, and 0.34 for smoothing, GPFA, and binning, respectively. Note that, for these offline analyses, the smoothing approach encompasses state-of-the-art brain-machine interface decoders such as the Kalman Filter, as detailed in (Willett et al. 2017) and Online Methods). We also determined performance as a function of population size by drawing random sub-samples from the neural population (**Fig. 2b**). LFADS using 25 (X velocity) or 50 (Y velocity) neurons outperformed the other techniques applied to the full population of 202 neurons.

We next tested whether the population dynamics inferred by LFADS on single trials exhibited dynamic features of motor cortical activity that have been identified previously by analyzing trial-averaged data. One such feature is the slow oscillations (~1-2.5 Hz) in motor cortical neural firing rates that accompany the transition from pre- to peri-movement activity in monkeys (Churchland et al. 2012) and humans (Pandarinath et al. 2015) that are consistent across the full range of movements being performed (**Fig. 3a**, monkey J showing consistent dynamics across 108 reach conditions of the “maze” dataset, and **Fig. 3c**, participant T5 showing consistent dynamics across 8 attempted movement conditions in a “center-out” task). These results were obtained by averaging the firing rate of each neuron across all the trials corresponding to a particular reach condition (condition averaging), and then applying a form of dimensionality reduction (jPCA). Although condition-averaging reveals the basic oscillatory dynamics, jPCA performed on single trials provides noisy and unstructured views of the neural trajectories (**Figs. 3b & 3d**). LFADS not only reproduces the previously-extracted oscillatory dynamics on a condition-averaged basis (**Figs. 3e & 3g**), it also clearly demonstrates, for the first time, the presence and consistency of oscillatory dynamics on single trials (**Fig. 3f**, monkey J, 2296 maze reaching trials, and **Fig. 3h**, participant T5, 114 trials of center-out movement attempts). The consistency of these trajectories at the single trial level is also clearly demonstrated in movies of the neural population state space trajectories over time (**Supp. Video 1**).

**Figure 3.**
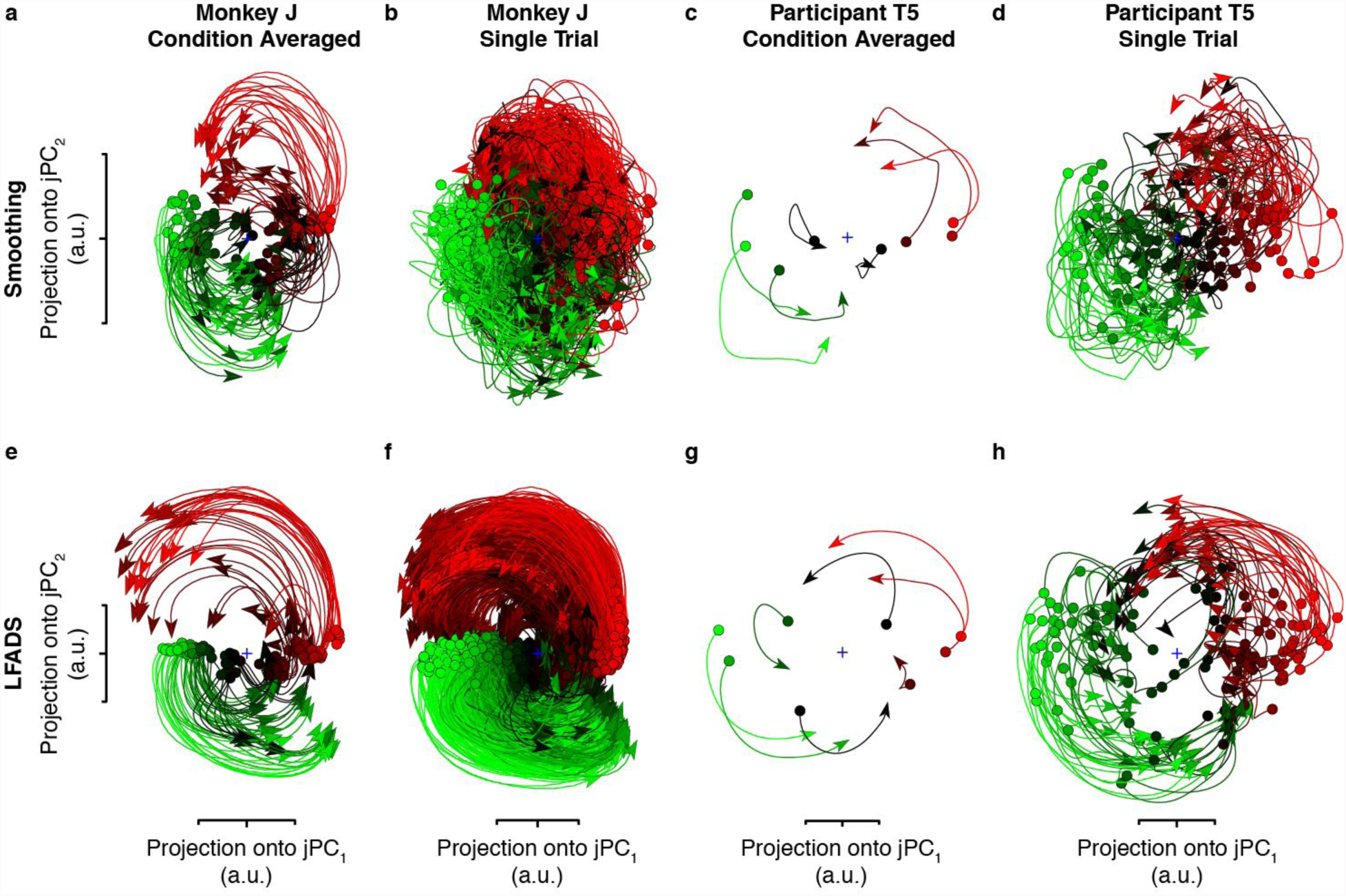
LFADS uncovers known slow dynamics in monkey and human motor cortical activity on a single-trial basis. (**a, c**) Slow oscillations of the neural population state accompany the transition between pre- and peri-movement activity, and have been previously described for monkey (Churchland et al., 2012) and human (Pandarinath et al., 2015) motor cortical activity. Each trace shows the neural population state trajectories for a single task condition (monkey: 108 reaching conditions; human: 8 intended movement directions). These dynamics were previously uncovered by averaging each neuron’s response across all trials of a given condition, and then projecting this condition-averaged activity into a low-dimensional plane using the jPCA technique. (**b, d**) When the same low-dimensional projection is applied to the single-trial data, dynamics are less clear due to the inherent noise of single-trial neural population activity. (**e, g**) De-noising the data via LFADS reveals the underlying dynamic structure at the condition average level for monkey and human. (**f, h**) Additionally, the same dynamic structure is now clearly present on individual trials (monkey: 2296 trials; human: 114 trials).

### LFADS rate oscillations and the local field potential

The previous analysis demonstrated that LFADS is capable of uncovering known slow oscillations in motor cortical firing rates on a single-trial basis. We also tested whether LFADS is capable of extracting dynamic features at faster timescales. A second known dynamic feature of motor cortical activity is the rhythmic spiking activity that often occurs during the pre-movement period in reaching tasks, typically phase-locked to accompanying oscillations in recorded LFPs [15-40 Hz; e.g., (Murthy and Fetz 1996; Donoghue et al. 1998)]. Detecting these dynamics on single trials requires capturing temporal structure in firing rates with sampling at the 7.5-20 ms timescale, a challenging problem given relatively infrequent spiking and trial-to-trial variability in motor behavior. We asked whether the oscillations evident in the inferred firing rates of **Fig. 1b** (seen in more detail in **Fig. 4a**), which were extracted without reference to LFPs, reflect this phenomenon. Indeed, fast oscillations in the inferred single-trial firing rates aligned well with LFPs and with structure apparent on single trials in unsorted, multi-unit spiking activity (i.e., threshold crossings; **Fig. 4a**).

**Figure 4.**
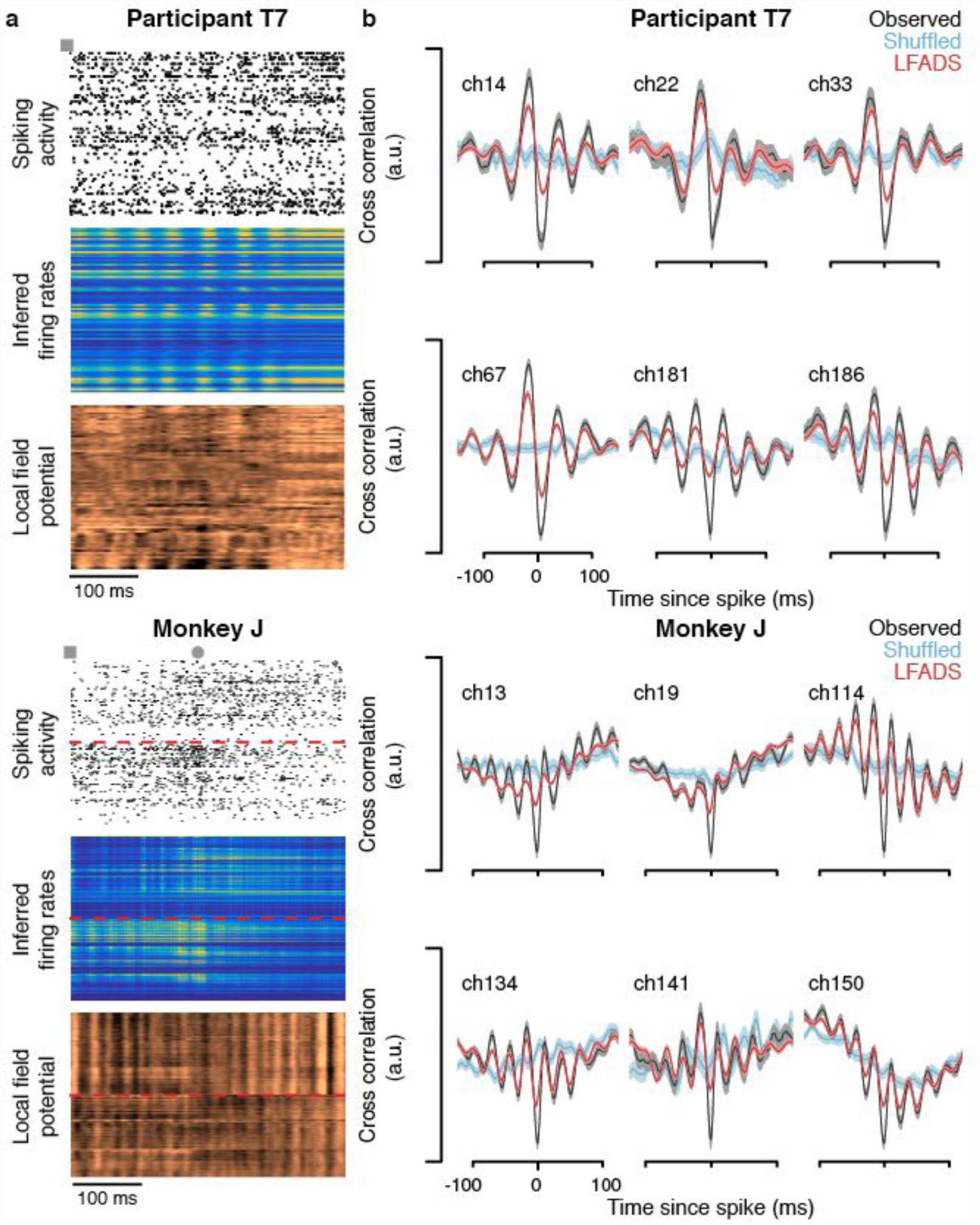
LFADS uncovers fast oscillatory structure in neural firing patterns. Previous work has demonstrated that spiking activity is often phase-locked to high-frequency oscillations in local field potentials (LFPs) prior to movement initiation (Murthy & Fetz, 1996; Donoghue et al., 1998). (**a**) Example single-trial neural activity recorded from human M1 and monkey M1/PMd. 400 ms of data are shown, beginning at the time of target presentation during an 8-target center-out-and-back movement paradigm. For T7, analyses were restricted to channels that showed significant modulation during movement attempts (78/192 channels). Dashed red lines overlaid on monkey data segregate the M1 array (upper halves) and PMd array (lower halves). Fast oscillations are evident in the spike rasters, the LFADS-inferred firing rates, and the recorded LFPs (LFPs were not used to train the LFADS models). Squares denote time of target onset. For Monkey J, where movement was measurable, circle denotes time of movement onset. (**b**) Cross-correlations between the observed spiking activity and the local field potentials recorded on the same electrode (black traces; mean ± s.e.m.) for several example channels that showed clear spike-LFP phase locking. Cross-correlations were computed for each channel on a single trial basis, and the resulting correlograms were averaged over all trials (participant T7: 142 trials; monkey J: 373 trials). LFP were first low-pass filtered (75 Hz cutoff frequency). Oscillations in these cross-correlations are indicative of consistent phase-locking between spikes and LFP. As expected, randomly shuffling the trial identity (i.e., correlating spikes from one trial with LFP from another) largely removed the fast, oscillatory components in the cross-correlograms (blue traces), demonstrating that the phase-locking is a single-trial phenomenon. Correlating the LFADS-inferred firing rates with recorded LFP (red traces) reveals that the inferred oscillatory structure in the neural firing rates corresponds with the recorded LFP in a similar fashion, despite LFADS having no access to the LFP signal.

Phase locking between spiking activity and LFPs in both monkey and human data is revealed by a cross-correlation analysis (**Fig. 4b**, black traces). Cross-correlations were first computed on a single-trial basis, using data from the first 250 ms (monkey) or first 300 ms (human) of each trial, and then averaged over trials. Averaging cannot be done before the cross-correlation is computed because this is a single-trial phenomenon; high-frequency oscillations in the cross-correlograms disappear when they are computed after shuffling trial identity (**Fig. 4b,** blue traces). As a result, correlations between condition-averaged firing rates and LFPs are almost certain to be averaged away.

The correlation between the firing rate oscillations revealed by LFADS on single trials and recorded LFPs is strikingly similar to the phase locking of the spiking activity with the LFP (**Fig. 4b**, red traces). This agreement is notable because no LFP data were used when training the LFADS model, and it provides further indication that the oscillations extracted by LFADS on single trials are a valid feature of the data.

### Stitching together data from multiple sessions

Thus far, we have demonstrated the application of LFADS to data recorded from a single neural population, but experiments are often performed across multiple sessions, with different populations of neurons recorded on each session, whether due to positional drift of a chronic array, or use of acute probes (e.g., v-probes) that are placed independently each session. Here, we ask if we can improve performance by using LFADS to stitch together data across multiple sessions, even when different sets of neurons are being recorded. LFADS provides a new ability to “stitch” such data together into a single dynamical systems model that can be used to analyze neural datasets collected over multiple days.

In an experimental setup where a subject is engaged in the same behavior across recording sessions and the same brain region is being recorded, it is reasonable that population samples composed of different neurons should participate in the same underlying dynamics, even if they are recorded at different times. LFADS is well suited to take advantage of this because of its two-step process of rate inference. LFADS can use the same generator network and dynamic factor readout matrices ***W^fac^*** across sessions, but employ different matrices ***W^rate^*** to infer firing rates for the different sets of neurons recorded during different sessions (**Fig. 5a**). Specifically, in this approach, the matrix ***W^fac^*** that computes the dynamic factors from the generator RNN is “learned” using data from all the sessions and, once learned, is held constant across all sessions. In contrast, separate ***W^rate^*** matrices, mapping from factors to firing rates, are learned from and applied to each session exclusively (see Online Methods for full details).

**Figure 5.**
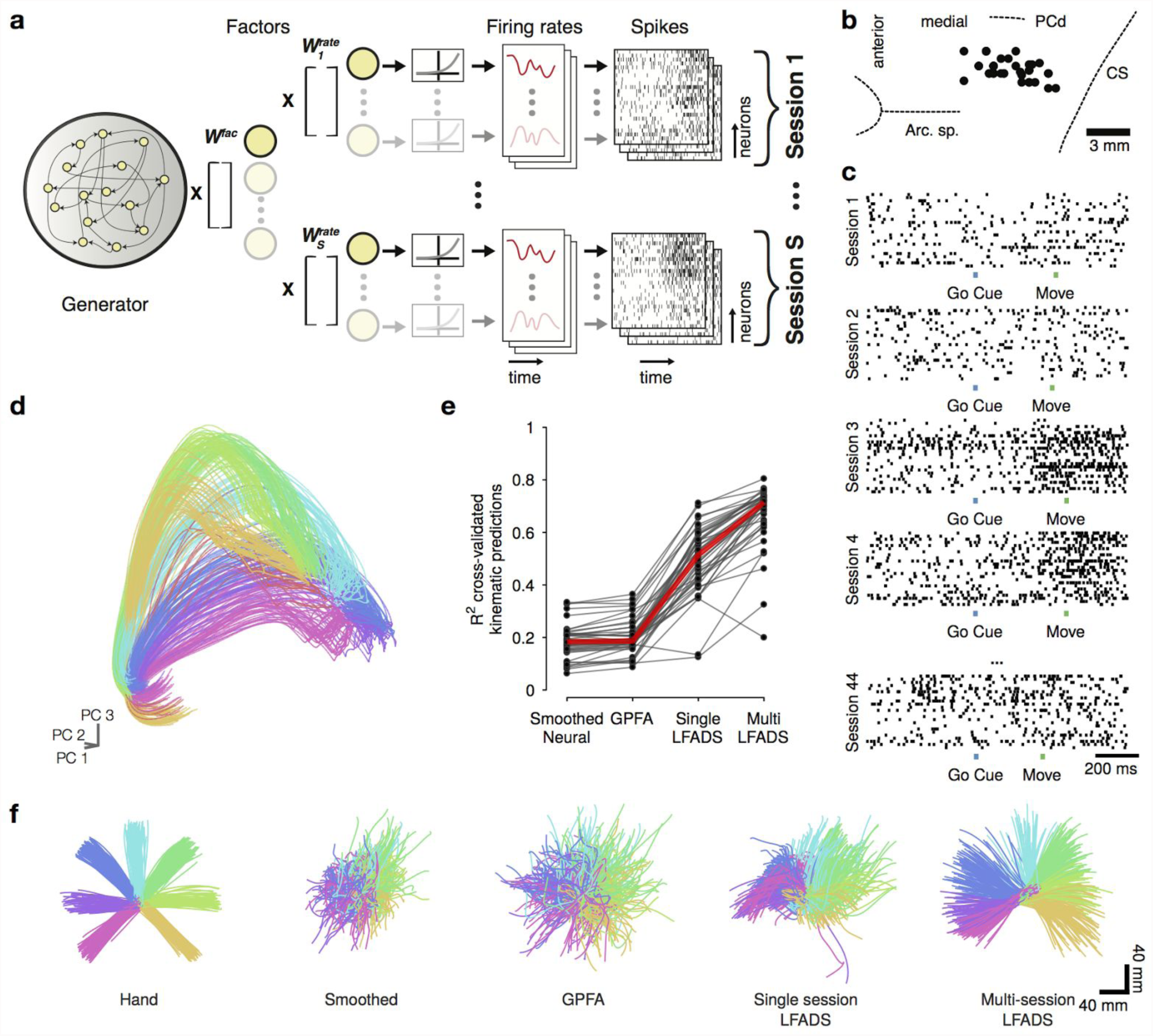
Using “dynamic neural stitching”, LFADS combines data from separately collected, non-overlapping recordings of the neural population by learning one consistent dynamical model. (**a**) The LFADS architecture was adapted to learn separate readouts from the dynamical generator for each v-probe recording session. I.e. a separate mapping from factors to firing rates 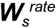 was learned for each session, while a single factor representation was used across sessions (i.e., only one readout matrix ***W****^fac^* is learned. 44 individual recording sessions were used. **(b)** Locations of linear electrode array penetrations in the precentral gyrus from which each dataset was collected. Dashed lines indicate approximate locations of nearby sulcal features based on stereotaxic locations. Arc. Sp.: arcuate spur, PCd: precentral dimple, CS: central sulcus. **(c)** Example single-trial rasters for nearly identical upwards reaches performed on a subset of 6 of the 44 recording sessions. Each raster has 24 rows corresponding to the 24 channels of the linear array, but the neurons recorded on each session are entirely distinct from each other. **(d)** After training, the multi-session stitched LFADS model produced consistent factor trajectories across recording sessions. Traces are factor trajectories for the multi-session stitched LFADS model projected into the top 3 principal component space. Each trajectory represents the single-trial factors time-series projected into the PCA subspace, while the color of the trajectory represents the reach direction. The spatial proximity of the reach trajectories for a given direction across the sessions (44 trajectories of each color) illustrates the consistency of the representation across sessions. **(e)** Using kinematic decoding (OLE), we observe that the factors of the multi-session stitched model are more informative about behavioral parameters (arm reaching kinematics) than the factors of LFADS models fit to each session individually. This is shown as a comparison of R^2^ values between arm kinematics and either smoothing neural data, GPFA, single-session or multi-session LFADS decodes. **(f)** Actual recorded hand position traces for center out reaching task, alongside kinematic decodes for a representative single session (32), for smoothed neural data, GPFA, single-session LFADS, and multi-session LFADS. Colors indicate reach direction.

We tested this approach using neural activity from monkey M1 and PMd during a center-out instructed-delay reaching task, recorded using linear multi-electrode arrays (monkey P; 24 channel V-probes, Plexon Inc.). We trained one stitched multi-session LFADS model on a combined dataset consisting of 44 recording sessions, which included 38 separate electrode penetration sites and spanned 162 days (**Fig. 5b** shows locations of the individual penetration sites in the precentral gyrus, and **Fig. 5c** shows sample recordings from 6 sessions). We then examined the condition-averaged factor trajectories traversed by the model for each recording session, for each reach direction. These trajectories are highly similar for a given reach direction, regardless of the recording session (**Fig. 5d**), a key indication that LFADS found a dynamical generator capable of producing a consistent set of factors. The consistency of the trajectories at the single trial level across the recording sessions is demonstrated in movies of the neural population state space trajectories over time (**Supp. Video 2**).

We then compared the multi-session LFADS model to 44 “single-session” models, each individually trained using data from a single session, using the same hyperparameter settings. We assessed the quality of the LFADS models by asking how informative the dynamic factors were in predicting behavioral observations, including reach kinematics and reaction time. In this case we decode from the dynamic factors because we are testing whether the representation across recording sessions is improved by LFADS, whereas inferred rates in both multi-session and single session models tend to be similar since, in a given session, LFADS almost always fits the observed population well. As expected from the previous analyses, the single-session LFADS models produced representations that were substantially more predictive of kinematics than Gaussian-smoothed neural data (mean improvement of 0.31 in R2; p < 10^−6^, Wilcoxon signed-rank test) or GPFA- smoothed neural data (mean improvement of 0.29 in R^2^, p < 10^−6^, Wilcoxon signed-rank test; **Fig. 5e),** indicating that LFADS identified useful dynamic representations even when operating on the limited observations from individual experimental sessions. Further, the multi-session LFADS model produced representations that were considerably better than even the single-session LFADS models, enabling kinematic predictions that greatly outperformed the single-session LFADS models (mean increase of 0.18 in R^2^, p < 10^−6^, Wilcoxon signed-rank test; **Fig. 5e, f**). We also predicted reaction time from LFADS factors using an unsupervised method [thresholding a condition-independent signal, see (Kaufman et al. 2016) and Online Methods]; again the stitched model significantly outperformed the single-day models (mean improvement in correlation coefficient between predicted and measured reaction times: 0.17; p < 10^−6^, Wilcoxon signed-rank test; **Supp. Fig. 5**).

### Inferring inputs to a neural circuit

Unmeasured input from unrecorded brain areas is a major source of ambiguity in studies of neural activity. Such inputs play an obviously important role if something unexpected occurs during a trial. The ability of LFADS to infer inputs ***u_t_*** (see Online Methods eqn. 15 for precise definition of ***u_t_*** and its role in the LFADS architecture) relies on the assumption that, if a powerful nonlinear dynamical system cannot generate a particular data set autonomously, it is likely that an external perturbation to the system occurred. Within the LFADS architecture, we extract this perturbation and identify it as an inferred input. Inferred inputs can be viewed as estimates of the inputs to the neural circuit being recorded (we outline caveats in the Discussion). At a minimum, inferred inputs are informative of the presence, type and timing of the unexpected perturbations during a trial (see **Supp. Figs. 6-8** for examples of inferring inputs to synthetic dynamical systems where the ground truth is known, e.g. the ‘integration to bound’ model in **Supp. Fig. 8**).

To test this approach, we analyzed data from a “Cursor Jump” task in which a monkey guided a cursor, controlled by the monkey’s hand position, to reach towards upward or downward targets (monkey J; see Online Methods). The target position (upward or downward) was shown to the monkey starting at the beginning of the trial (target onset). In “unperturbed” trials (75%), the cursor consistently tracked the position of the monkey’s hand, and the monkey made straight upward or downward reaching movements to acquire targets. On 25% of the trials (“perturbed” trials), unpredictable shifts of 6 cm to the left or right between cursor and hand position forced the monkey to make corrective movements to acquire the target (**Fig. 6a**). We applied LFADS to spiking activity from multi-electrode arrays implanted in M1 and PMd (**Fig. 6b**), allowing four inferred inputs. We analyzed the first 800 ms of each trial (beginning at target onset; most jumps occurred 350-550 ms after target onset).

**Figure 6.**
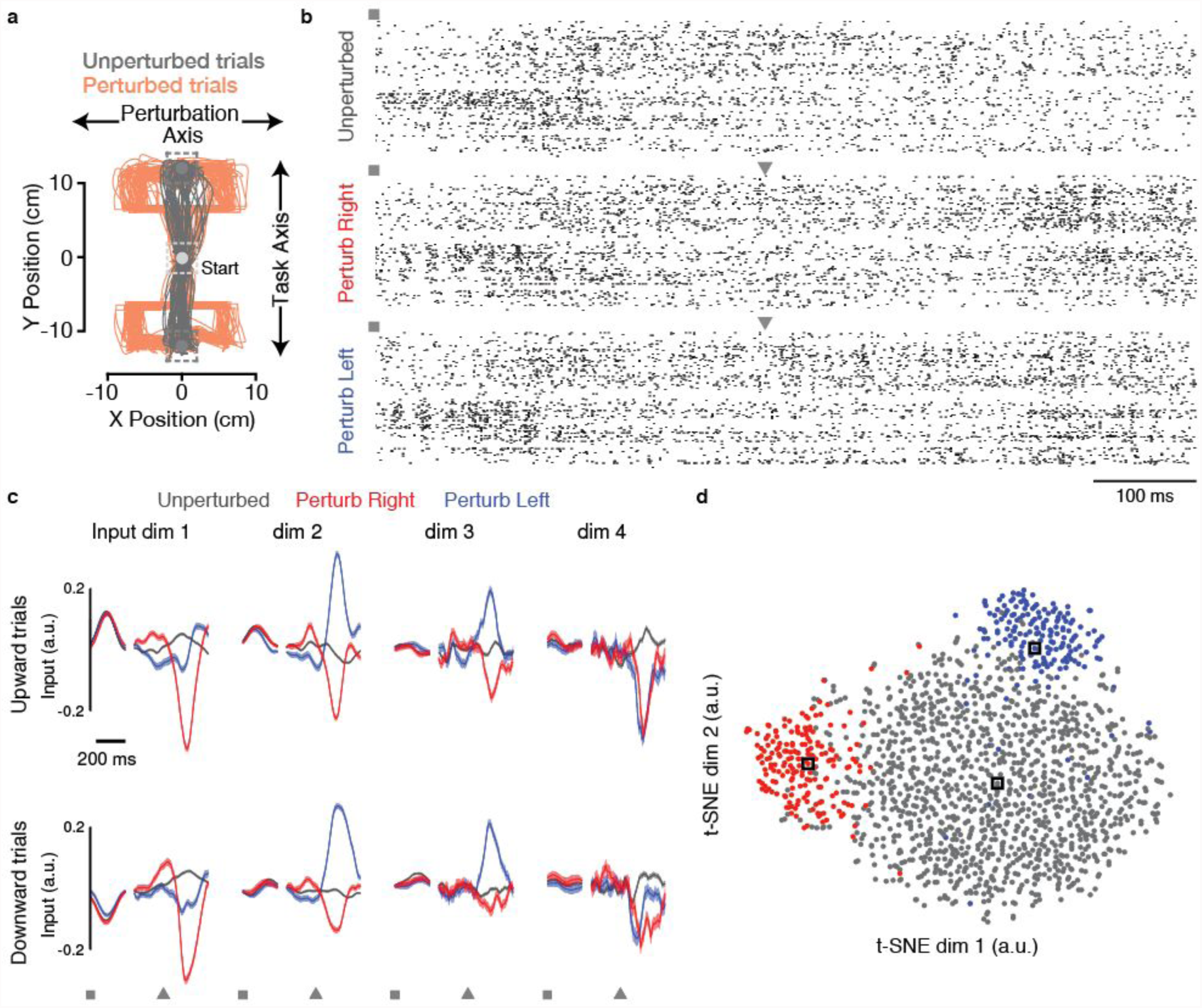
LFADS uncovers the presence, identity and timing of unexpected perturbations in the “Cursor Jump” task. (**a**) The position of a monkey’s hand was linked to the position of an on-screen cursor, and the monkey made reaching movements to steer the cursor toward upward or downward targets. In unperturbed trials (grey traces), the monkey made straight reaches to the target. In perturbed trials (orange traces), the cursor’s position was offset to the left or right during the course of the reaching movement, and the monkey made corrective movements to acquire the target. (**b**) Spiking activity from M1/PMd arrays during three example reach trials towards downward targets for the unperturbed (top), perturb right (middle), and perturb left (bottom) conditions. Squares denote time of target onset, and triangles denote the time of an unexpected perturbation. (**c**) LFADS was allowed 4 inferred inputs to model the neural activity. For presentation, two trial alignments were used prior to averaging: the initial portion of the trials was aligned to the time of target onset, while the latter portion of the trials was aligned by perturbation time (or, for unperturbed trials, the time at which a perturbation would have occurred based on the cursor’s trajectory). The gap in the traces denotes the break in alignment. Inferred input values were averaged across trials for upward (top) and downward (bottom) trials (mean ± s.e.m. is shown, grey: unperturbed trials, blue: perturb left trials, red: perturb right trials). Around the time of target onset, the identity of the target (up vs. down) is modeled by the inputs (e.g., dimension 1). Around the time of the perturbation, LFADS used specific inferred input patterns to model each perturbation type (e.g., dimensions 1 & 2). Input traces were smoothed with a causal Gaussian filter (20 ms s.d.). (**d**) The single-trial input patterns around the time of perturbation (all downward trials) were projected into a low-dimensional space using t-SNE. The clustering of the three perturbation types (unperturbed, left perturbation, or right perturbation) is highly distinguishable, implying the single-trial inferred inputs are separable based on perturbation type. Black boxes denote locations in t-SNE space for the example trials shown in panel **b**.

LFADS inferred inputs to model information flow into the generator with timing that was consistent with the trial structure. Prior to the trial, the monkey had no information about the target position, which was cued at the beginning of the trial (target onset). Around this time, the inferred inputs are distinct with respect to target position (**Fig. 6c**, e.g. Input dim 1, comparing inputs inferred for Upward trials vs. Downward trials), but are not distinct with respect to perturbation type (i.e., red, blue, and grey traces are overlapping), as the perturbation was not presented to the monkey until later in the trial. In contrast, around the time of perturbation, LFADS inferred input patterns that were different for right- and left-shift perturbed trials and for unperturbed trials (**Fig. 6c**, red, blue, and grey traces, e.g. Input dim 2). Furthermore, the timing of these inputs is well aligned to the time of the perturbations (which were variable), and the perturbation direction specificity of these inputs were similar across downward and upward reaches (**Fig. 6c**, top and bottom panels). The described trend in the inferred inputs were also visible at the single trial level (**Supp. Fig. 9**). We note that the exact shape of the inferred inputs may not resemble physiological signals. In addition, the timing of the inputs is not required to be causal relative to the timing of the perturbations (see Discussion). Nevertheless, this example demonstrates the ability of LFADS to predict, on average, the presence, identity and timing of inputs to motor cortex related to task perturbations.

Finally, to test whether these inputs contained information about the identity of the perturbation on single trials, we applied a widely used nonlinear dimensionality reduction technique, t-distributed stochastic neighborhood embedding (t-SNE), to the inferred single-trial inputs around the time of the perturbation (**Fig. 6d**). t-SNE revealed that the inferred inputs cluster according to the identity of the perturbation on a single-trial basis.

## Discussion

The ability to record from large ensembles of neurons has inspired a shift from emphasizing the properties of individual neurons and their responses to exploring the dynamics of large neural populations [see e.g. (Yuste 2015)]. LFADS aids in this process by inferring the underlying dynamics and states of a neural population on single trials. In particular, LFADS constructs a de-noised time-dependent firing rate for each neuron on each trial. To obtain a reduced description of the data, there is no need to pass these firing rates through an additional dimensionality reduction step because the dynamic factors generated by LFADS *are* a reduced description. Furthermore, there is no need to construct a network model that can account for these rates because this is provided by the generator RNN.

The population state inferred by LFADS is likely to be considerably more informative about an animal’s behavior than raw single-trial observations, even when a large population of neurons is observed (**Fig. 2**). This is because the population state inferred by LFADS is informed by its model of the system’s underlying dynamics, which leverages all the observed data. Further, LFADS attempts to infer inputs to a brain region, and can also stitch together datasets from different recording sessions. We believe the interaction of these capabilities will provide a significant step forward in understanding the role of dynamics in driving computations in many behavioral paradigms and brain areas. For example, modeling complex dynamics and inputs to a brain area could be a powerful tool for understanding integration of evidence in complex decision-making tasks, for understanding how motor areas set up dynamic trajectories and integrate feedback/perturbations to guide corrective movements, or for understanding flexible, context-dependent communication between different brain areas.

There are a number of differences between LFADS and other neural population analysis techniques (Yu et al. 2009; Petreska et al. 2011; Macke et al. 2011; Kao et al. 2015; Aghagolzadeh and Truccolo 2014; Zhao and Park 2017; Y. Gao et al. 2016; Linderman et al. 2017), but its primary innovations are the use of nonlinear RNNs for the dynamics generator, the ability to infer inputs describing features of the data that cannot be fit by the dynamics model, and the ability to stitch together different datasets. As demonstrated using synthetic data, where ground truth dynamics are known (**Supp. Materials**), LFADS significantly outperforms other methods (see Online Methods for a more in-depth discussion of related work in the broader machine learning context).

A vexing challenge in neuroscience is distinguishing the role of dynamics internal to a neural circuit from the effects of unmeasured time-varying input from other brain regions. In the LFADS approach, inputs are inferred to account for features of the data that the learned autonomous RNN dynamics cannot, by themselves, explain. In this sense, inputs inferred by LFADS are similar to innovations in Kalman filtering. Our results on the Cursor Jump task (**Fig. 6**) show that LFADS can infer input fluctuations that coincide in time with experimentally induced perturbations, and also that the inferred inputs on a given trial are informative about the existence and direction of the perturbation on that trial.

Although the nature of the inputs inferred by LFADS is informative about the presence and identity of perturbations, caution should be used when interpreting the precise shape and timing of these inputs. In addition to reflecting actual inputs that a neural circuit receives, LFADS-inferred inputs may reflect model mismatch (e.g. a biological spiking neural network vs. an artificial RNN) and measurement noise. Furthermore, there is no constraint requiring the shape of the inferred input to conform to physiological processes, and the transformation from the inferred input to the predicted neural activity is nonlinear and likely to be complex. Also, the timing of the inferred inputs may be shifted relative to the timing of the perturbations they describe. Finally, due to the bidirectional encoders used by LFADS, the generator has access to the entire data sequence being modeled, and there is no constraint forcing the inputs to be causal with respect to the task perturbation. Caveats aside, both the presence, timing, and qualitative shape of the inferred input in the Cursor Jump task (as well as in two synthetic examples) are reasonable, providing evidence that LFADS inferred inputs are likely to be useful for thinking about neural computations, e.g. by allowing computational models to disambiguate between internal dynamics from input driven dynamics.

LFADS provides a new ability to combine data from separate recording sessions involving different recorded neurons into a single dynamical systems model. The models LFADS constructs from such stitched data are likely to be more accurate than models extracted from any single session alone (**Fig. 5**). This argues that consistent dynamics can describe the activity of large functionally connected groups of neurons, even in recordings that span many different electrode penetrations from separate sessions across several months. In this study we used the stitching approach in a case where a relatively small number of neurons was simultaneously recorded in each session. However, if the identity of individual neurons changes or is difficult to track across sessions, stitching may prove useful even in cases where a large number of neurons is simultaneously recorded in each session, by providing a natural method to relate their activity across sessions. It is also possible that the stitching approach could be applied to utilize data across many subjects to further improve data-driven modeling of neural circuits. For example, under the assumption that motor cortical dynamics are similar across subjects, the same generator could be trained using many subjects’ data, while the differences in subjects’ data could be modeled using individualized output matrices or feedforward neural networks.

The single-trial neural population state inferred by LFADS facilitates more accurate decoding of external variables such as kinematic intention and reaction time. This holds potential for improving the performance of therapeutic neurotechnologies such as brain-machine interfaces (BMIs). Additionally, the ability of stitching to infer neural population state estimates that are invariant to the particular population being recorded may further improve BMI stability. To leverage these innovations for BMI applications, two changes would be required: first, a real-time implementation of LFADS would have to be developed. This is feasible because, while *training* the LFADS model parameters is a computationally demanding task, *running* the LFADS model involves a series of linear transformations and static nonlinearities, which are on the same order of computational complexity as existing BMI implementations that run in real-time (Sussillo et al. 2012; Gilja et al. 2015; Sussillo et al. 2016; Pandarinath et al. 2017). Second, the LFADS architecture must be modified to be causal, i.e., the current neural state estimate must depend only on previously observed data. Beyond these two requirements, BMIs may be additionally enhanced via neural stitching, which could be harnessed for real-time applications by training most of the LFADS model parameters based on historical data, and simply updating the weights of the input and output adaptor matrices for the currently observed neural data. Alternatively, it may be possible to replace the these adaptor matrices with nonlinear architectures that adapt to the currently-observed data, as in (Sussillo et al. 2016).

There are multiple extensions and future directions to explore. First, an output model relevant to calcium imaging (replacing the Poisson spiking model) would allow the approach to be extended to calcium imaging data. Second, the LFADS generator could be strengthened by stacking recurrent layers or using a deep feedforward network architecture. Another extension would be to optimize the number of inferred inputs and dynamics factors along with the other model parameters, instead of fixing them to a specific value. Finally, to model areas which are considered to be more input-driven (e.g., early visual processing), an LFADS-like model that relies more on feedforward computation but still has recurrence may facilitate more accurate modeling of phenomena such as short-term adaptation.

Finally, by testing whether recordings from two data sets can be fit by the same generator RNN and factor-extraction matrices but different rate-extraction matrices, LFADS could be used to check whether the dynamics in two brain regions (or in two subjects) are consistent. For example, LFADS could be used to determine whether two brain regions are fulfilling similar or different functions, depending on whether stitching the two data sets provides a good or a bad fit, respectively. Given that the expansion from single-neuron to multi-neuron recording is now being followed by a shift from single- to multi-regional studies, LFADS may find widespread use.

## Acknowledgments

We would like to thank John P. Cunningham, Laurent Dinh, and Jascha Sohl-Dickstein for extensive conversation. We also thank Christine Blabe and Paul Nuyujukian for assistance with research sessions with participant T5, Emad Eskandar for array implantation with participant T7, and Brittany Sorice and Anish Sarma for assistance with research sessions with participant T7. R.J. participated in this work while at Google, Inc. L.F.A.’s research was supported by US National Institutes of Health grant MH093338, the Gatsby Charitable Foundation through the Gatsby Initiative in Brain Circuitry at Columbia University, the Simons Foundation, the Swartz Foundation, the Harold and Leila Y. Mathers Foundation, and the Kavli Institute for Brain Science at Columbia University. C.P. was supported by a postdoctoral fellowship from the Craig H. Neilsen Foundation for spinal cord injury research and the Stanford Dean’s Fellowship. S.D.S. was supported by the ALS Association’s Milton Safenowitz Postdoctoral Fellowship. K.V.S.’s research was supported by an NIH Director’s Pioneer Award, an NIH-NINDS T-RO1, NIH-NINDS R01NS066311, DARPA REPAIR, and the Simons Foundation. J.M.H.’s research was supported by NIH-NIDCD R01DC014034. K.V.S. and J.M.H.’s research was supported by Stanford BioX-NeuroVentures, Stanford Institute for Neuro-Innovation and Translational Neuroscience, Garlick Foundation and Reeve Foundation. L.R.H’s research was supported by NIH-NIDCD R01DC009899, Rehabilitation Research and Development Service, Department of Veterans Affairs (B6453R), MGH-Deane Institute for Integrated Research on Atrial Fibrillation and Stroke; Executive Committee on Research, Massachusetts General Hospital.

The content is solely the responsibility of the authors and does not necessarily represent the official views of the National Institutes of Health, the Department of Veterans Affairs, or the United States Government. BrainGate CAUTION: Investigational Device. Limited by Federal Law to Investigational Use.

## 1 The LFADS Model

A TensorFlow reference implementation of the LFADS model will soon be made available.

### 1.1 The variational auto-encoder

The LFADS model is an instantiation of a variational auto-encoder (VAE) [19, 29] extended to sequences, as in [13] or [21]. The VAE consists of two components, a decoder (also called a generator) and an encoder. The generator assumes that data, denoted by **x**, arise from a random process that depends on a vector of stochastic latent variables **z**, samples of which are drawn from a prior distribution *P*(**z**). Simulated data points are then drawn from a conditional probability distribution, *P*(**x**|**z**) (we have suppressed notation reflecting the dependence on parameters of this and the other distributions we discuss).

The VAE encoder transforms actual data vectors, **x**, into a conditional distribution over **z**, *Q*(**z**|**x**). *Q*(**z**|**x**) is a trainable approximation of the posterior distribution of the generator, *Q*(**z**|**x**) ≈ *P*(**z**|**x**) = *P*(**x**|**z**)*P*(**z**)*/P*(**x**). *Q*(**z**|**x**) can also be thought of as an encoder from the data to a data-specific latent code **z**, which can be decoded using the generator (decoder). Hence the auto-encoder; the encoder *Q* maps the actual data to a latent stochastic “code”, and the decoder *P* maps the latent code back to an approximation of the data. Specifically, when the two parts of the VAE are combined, a particular data point is selected and an associated latent code, 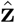 (we use 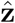 to denote a sample of the stochastic variable **z**) is drawn from *Q*(**z**|**x**). A data sample is then drawn from 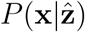, on the basis of the sampled latent variable. If the VAE has been constructed properly, 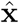 should resemble the original data point **x**.

The loss function that is minimized to construct the VAE involves minimizing the Kullback-Leibler divergence between the encoding distribution *Q*(**z**|**x**) and the prior distribution of the generator, *P*(**z**), over all data points. In the VAE framework *P*(**z**) is typically defined as a Gaussian prior whose parameters are independent of the data, and that is also the case here. The rationale is that the even a simple distribution, such as a Gaussian, can be transformed into a complex distribution by passing samples of the Gaussian distribution through a powerful nonlinear function. If training is successful, *Q*(**z**|**x**) and *P*(**z**) should converge and, in the end, statistically accurate generative samples of the data can be created by running the generator model seeded with samples from *P*(**z**), i.e. accurate samples of the data can be generated from white noise.

We now translate this general description of the VAE into the specific LFADS implementation aimed at high-dimensional, simultaneously recorded neural spike trains. Borrowing some notation from [13], we denote an affine transformation (**v** = **W u** + **b**) from a vector-valued variable **u** to a vector-valued variable **v** as **v** = **W**(**u**), we use [·, ·] to represent vector concatenation, and we denote a temporal update of a recurrent neural network receiving an input as state*_t_* = RNN^a^(state*_t_*_−1_, input*_t_*), for an RNN named’a’. It is understood that if there are two networks modules, such as RNNs, with different names, e.g. RNN^a^(.,.) and RNN^b^(.,.), these network modules do not share parameters.

### 1.2 LFADS Generator

The neural data we consider, **x**_1:*T*_, consists of spike trains from *D* recorded neurons. Our reference implementation of LFADS also supports continuous Gaussian distributed data, but as this is not central to the main application, we focus exclusively on spike trains in what follows. Each instance of a vector **x**_1:*T*_ is referred to as a trial, and trials may be grouped by experimental conditions, such as stimulus or response types. The data may also include an additional set of observed variables, **a**_1:*T*_, that may refer to stimuli being presented or other experimental features of relevance, such as kinematics. Unlike **x**_1:*T*_, the data described by **a**_1:*T*_ is not itself being modeled, but it may provide important conditioning information relevant to the modeling of **x**_1:*T*_. This introduces a slight complication: we must distinguish between the complete data set, {**x**_1:*T*_, **a**_1:*T*_} and the part of the data set being modeled, **x**_1:*T*_. The conditional distribution of the generator, *P*(**x**|**z**), is only over **x**, whereas the approximate posterior distribution, *Q*(**z**|**x**, **a**), depends on both types of data.

LFADS assumes that the observed spikes described by **x**_1:*T*_ are samples from a Poisson process with underlying rates **r**_1:*T*_. Based on the dynamical systems hypothesis outlined in the introduction of the main text, the goal of LFADS is to infer a reduced set of latent dynamic variables, **f**_1:*T*_, of dimension *F*, from which the firing rates can be constructed. The rates are determined from the factors by an affine transformation followed by an exponential nonlinearity, **r**_1:*T*_ = exp(**W**^rate^(**f**_1:*T*_)). Note that exp(·) is the inverse canonical link function for the Poisson distribution, making it a natural choice to keep the Poisson rate variable positive. The choice of a low-d representation for the factors is based on the observation that the intrinsic dimensionality of neural recordings tends to be far lower than the number of neurons recorded, e.g. [6, 16, 24], and see [8] for a more complete discussion.

The factors are generated by a recurrent nonlinear neural network and are characterized by an affine transformation of its state vector, **f**_1:*T*_ = **W**^fac^(**g**_1:*T*_), with **g***_t_* of dimension *N*. Running the network requires an initial condition **g**_0_, which is drawn from a prior distribution 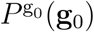. Thus, **g**_0_ is an element of the set of the stochastic latent variables **z** discussed above.

There are different options for sources of time-dependent input to the recurrent generator network. First, as in some of the examples to follow, the network may receive no input at all. Second, it may receive the information contained in the non-modeled part of the data, **a**_1:*T*_, in the form of a network input. Instead, as a third option, we introduce an inferred input **u**_1:*T*_. When an inferred input is included, the set of stochastic latent variables is expanded to include it, **z** = {**g**_0_, **u**_1:*T*_}. At each time step, **u***_t_* is drawn from a prior distribution *P*^u^(**u***_t_*|**u***_t_*_−1_) that is auto-regressive, with 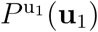 defining the distribution over **u**_1_. (see section 1.7).

The LFADS generator with inferred input is thus described by the following procedure and equations. First an initial condition for the generator is sampled from the prior on **g**_0_

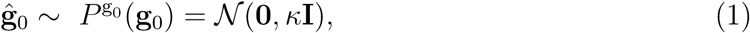

with *κ* a hyperparameter. At each time step *t* = 1,…, *T*, an inferred input, **û***_t_*, is sampled from its prior and fed into the network, and the network is evolved forward in time,

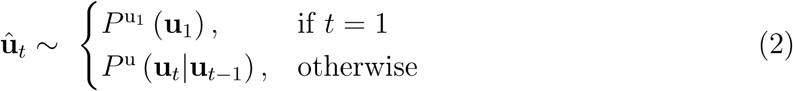

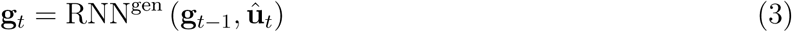

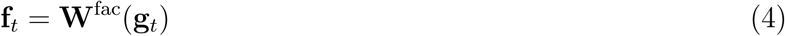

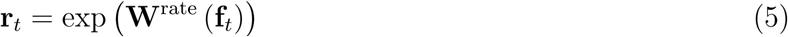

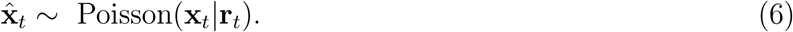

Here “Poisson” indicates that each component of the spike vector **x***_t_* is generated by an independent Poisson process at a rate given by the corresponding component of the rate vector **r***_t_*. The prior for both **g**_0_ and **u**_1_ are diagonal Gaussian distributions. The prior for **u***_t_* with *t* > 1 is an auto-regressive Gaussian prior, with a learnable autocorrelation time and process variance (see section 1.7 for more details). We chose the Gated Recurrent Unit (GRU) [4] as our recurrent function for all the networks we use (see section 1.6 for equations), including RNN^gen^. We have not included the observed data **a** in the generator model defined above, but this can be done simply by including **a***_t_* as an additional input to the recurrent network in equation 3. Note that doing so will make the generation process necessarily dependent on including an observed input. The generator model is illustrated in **Methods Fig. 1**. This diagram and the above equations implement the conditional distribution *P*(**x**|**z**) = *P*(**x**|{**g**_0_, **u**_1:*T*_}) of the VAE decoder framework.

**Figure 1:**
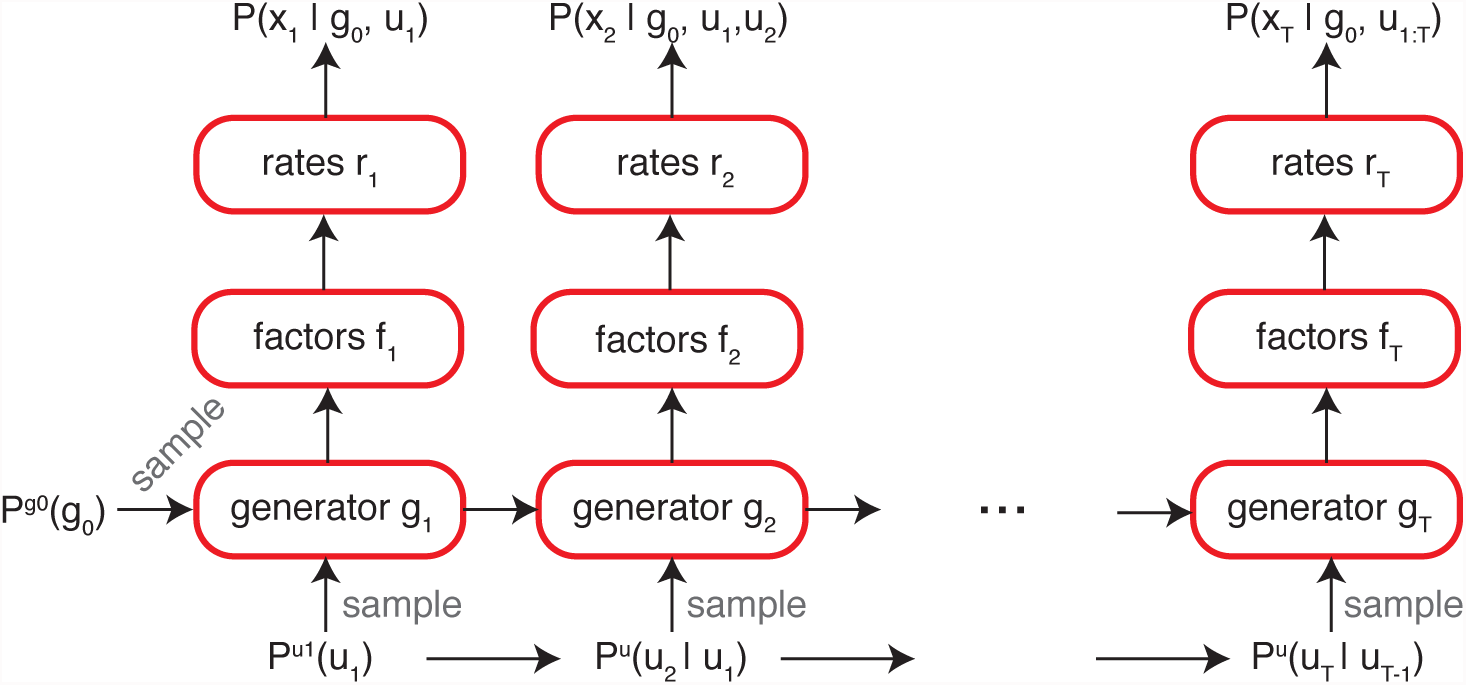
The LFADS generator. The generative LFADS model is a recurrent network with a feed-forward readout. The generator takes a sampled initial condition, 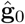 and a sampled inferred input, 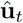, at each time step, and iterates forward. At each time step the temporal factors, **f***_t_*, and the rates, **r***_t_* are generated in a feed-forward manner from **g***_t_*. Spikes are generated from a Poisson process, 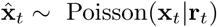. The initial condition and input at time step 1 are sampled from diagonal Gaussian distributions with zero mean and fixed chosen variance. Otherwise, the inputs are sampled from a Gaussian auto-regressive prior.

### 1.3 LFADS Encoder

The approximate posterior distribution for LFADS is the product of two conditional distributions, one for **g**_0_ and one for **u***_t_*. Both of these distributions are Gaussian with means and diagonal covariance matrices determined by the outputs of the encoder or controller RNNs (see **Methods Fig. 2** and below). We begin by describing the network that defines 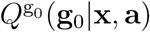. Its mean and variance are given in terms of a vector **E**^gen^ by

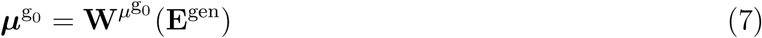

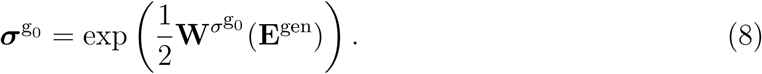

**Figure 2:**
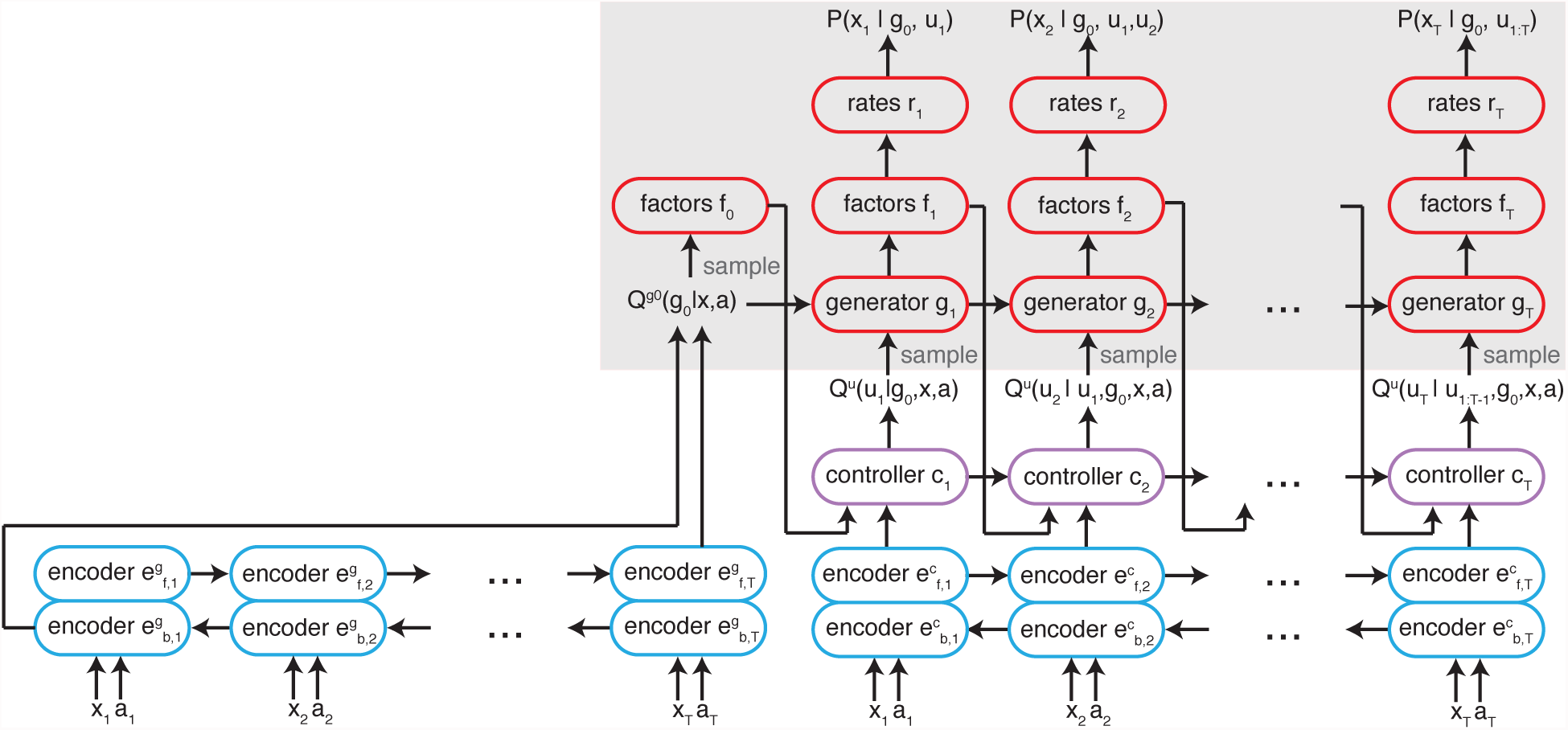
The full LFADS model for inference. The generator / decoder portion highlighted with a gray background and is colored red, the encoder portion is colored blue and the controller, purple. To infer the latent dynamics from the recorded neural spike trains **x**_1:*T*_ and conditioning data **a**_1:*T*_, initial conditions for the controller and generator networks are encoded from inputs. In the case of the generator, the initial condition 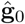 is drawn from an approximate posterior 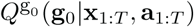 that receives an encoding of the input, **E**^gen^ (in this figure, for compactness, we use **x** and **a** to denote **x**_1:*T*_ and **a**_1:*T*_). The low-dimensional factors at *t* = 0, **f**_0_, are computed from 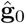. The controller then propagates one step forward in time, receiving the sample factors **f**_0_ as well as bidirectionally encoded inputs 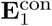 computed from **x**_1:*T*_, **a**_1:*T*_. The controller produces, through an approximate posterior *Q*^u^(**u**_1_|**g**_0_,**x**_1:*T*_,**a**_1:*T*_), a sampled inferred input 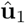 that is fed into the generator network. The generator network then produces {**g**_1_, **f**_1_, **r**_1_}, with **f**_1_ the factors, and **r**_1_ the Poisson rates at *t* = 1. The process continues iteratively so, at time step *t*, the generator network receives **g***_t_*_−1_ and 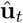 sampled from ***Q*^u^(u*_t_*|u_1:*t*−1_, g_0_, x_1:*T*_, a_1:*T*_**). The job of the controller is to produce a nonzero inferred input only when the generator network is incapable of accounting for the data autonomously. Although the controller is technically part of the encoder, it is run in a forward manner along with the decoder.

**E**^gen^ is obtained by running two recurrent networks over the data, bidirectionally. One RNN runs forward (from *t* = 1 to *t* = *T*) in time and the other RNN runs backwards (from *t* = *T* to *t* = 1),

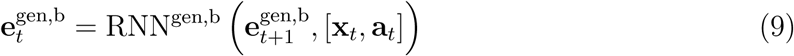

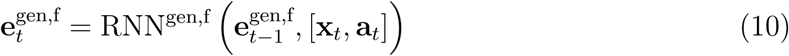

with 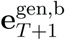 and 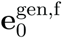 learnable biases. Once this is done, **E**^gen^ is the concatenation

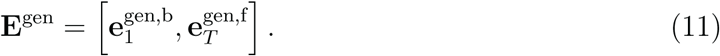

Running the encoding network both forward and backward in time allows **E**^gen^ to reflect the entire time history of the data **x**_1:*T*_ and **a**_1:*T*_. Finally, we sample initial conditions 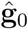 according to the following distribution

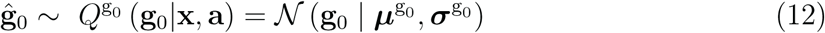

for a normal distribution with mean 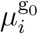 and standard deviation 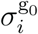 for the *i^th^* element of **g**_0_.

The approximate posterior distribution for **u***_t_* is defined in a more complex way that involves both a second set of forward-backward encoder RNNs and another RNN called the controller. The forward and backward encoder RNNs provide the input to the controller RNN, and are defined at time *t* with state variables 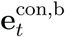 and 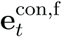 that are defined by equations identical to 9 and 10 (although with different trainable network parameters). Finally, the time-dependent input to the controller RNN is defined as

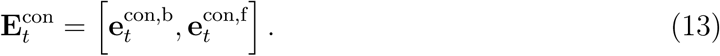

Rather than feeding directly into a Gaussian distribution, this variable is passed through the controller RNN, which runs forward in time with the generator RNN and also receives the latent dynamic factor, **f***_t_*_−1_ as input,

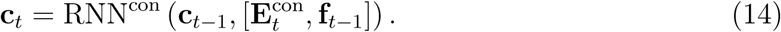

Thus, the controller is privy to the information about **x**_1:*T*_ and **a**_1:*T*_ encoded in the variable 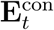, and it receives information about what the generator network is producing through the latent dynamic factor **f***_t_*_−1_. It is necessary for the controller to receive the factors so that it can correctly decide when to intervene in the generation process. Because **f***_t_*_−1_ depends on both **g**_0_ and **u**_1:*t*−1_, these stochastic variables are included in the conditional dependence of the approximate posterior distribution *Q*^u^(**u***_t_*|**u**_1:*t*−1_, **g**_0_,**x**_1:*T*_, **a**_1:*T*_). The initial state of the controller network, **c**_0_, is defined as a trainable bias initialized to the 0 vector.

Finally, the inferred input, **u***_t_*, at each time, is a stochastic variable drawn from a diagonal Gaussian distribution with mean and log-variance given by an affine transformation of the controller network state, **c***_t_*,

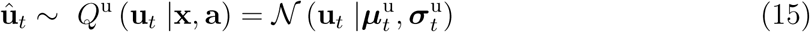

with

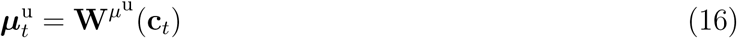

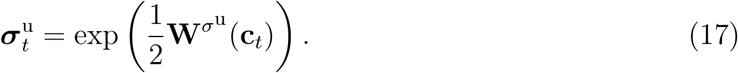

We control the information flow out of the controller and into the generator by applying a regularizer on **u***_t_* (a KL divergence term, described in Sections 1.5 and 1.9), and also by explicitly limiting the dimensionality of **u***_t_*, the latter of which is controlled by a hyperparameter.

### 1.4 The full LFADS inference model

The full LFADS model (**Methods Fig. 2**) is run in the following way. First, a data trial is chosen, the initial condition and inferred input encoders are run, and an initial condition is sampled from the approximate posterior, 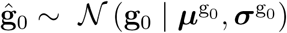. Then, for each time step from 1 to *T*, the generator is updated, as well as the factors and rates, according to

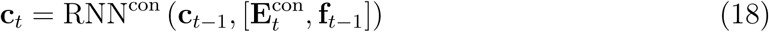

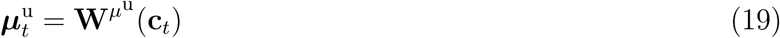

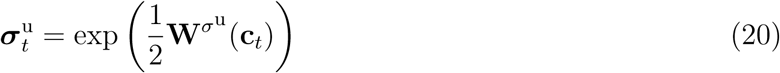

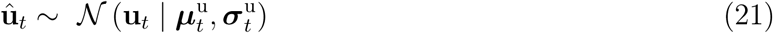

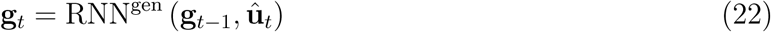

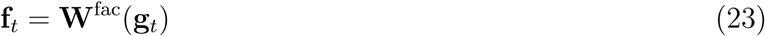

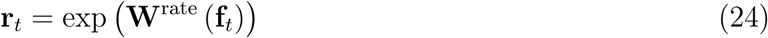

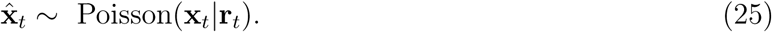

After training, the full model can be run, starting with any single trial or a set of trials corresponding to a particular experimental condition to determine the associated dynamic factors, firing rates and inferred inputs for that trial or condition. This is done by averaging over several runs to marginalize over the stochastic variables **g**_0_ and **u**_1:*T*_. Typically, equation 25 is not executed, unless one explicitly desires to generate spikes.

### 1.5 The loss function

To optimize our model, we would like to maximize the log likelihood of the data, 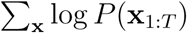, marginalizing over all latent variables. For reasons of intractability, the VAE framework is based on maximizing a variational lower bound, 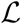, on the marginal data log-likelihood,

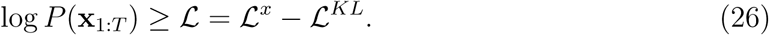

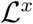 is the log-likelihood of the reconstruction of the data, given the inferred firing rates, and 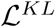 is a non-negative penalty that restricts the approximate posterior distributions from deviating too far from the (uninformative) prior distribution. 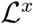 and 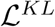 are then defined as

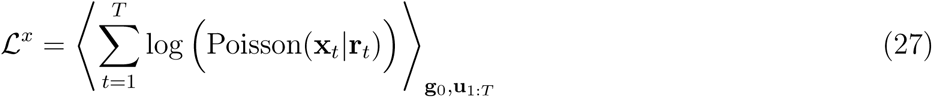

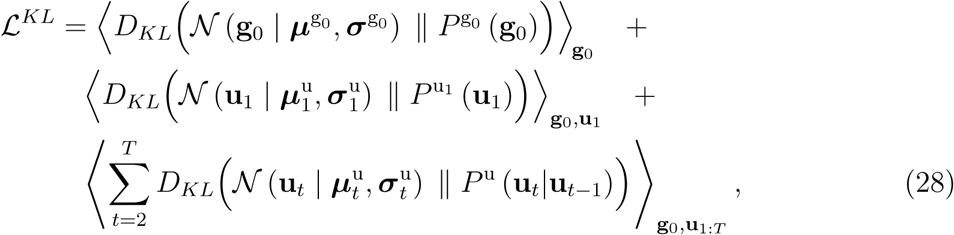

where the brackets denote marginalizations over the sub-scripted variables. Evaluating the *T* +1 KL terms can be done analytically for the Gaussian distributions we use; the formulae are found in Appendix B of [19]. We minimize the negative bound, 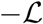, using the reparameterization trick for Gaussian distributions to back-propagate low-variance, unbiased gradient estimates [19]. These gradients are used to train the system in an end-to-end fashion, as is typically done in deterministic settings.

### 1.6 GRU equations

For clarity, we use the common variable symbols associated with the GRU, with the understanding that the variables represented here by these symbols are not the same variables as those in the general LFADS model description. For **x***_t_* the input and **h***_t_* the hidden state at time *t*, the GRU update equation, **h***_t_* = GRU(**x***_t_*, **h***_t_*_−1_), is defined as

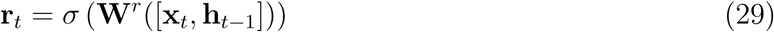

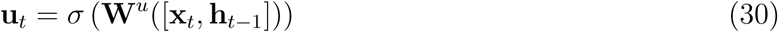

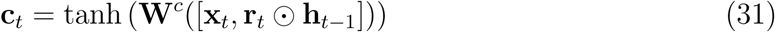

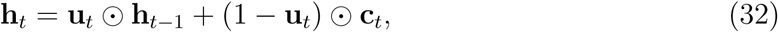

with ⊙ denoting element-wise multiplication and *σ* denoting the logistic function.

### 1.7 Autogressive prior for inferred input

A zero-mean auto-regressive process with one time lag (AR(1)) is defined by

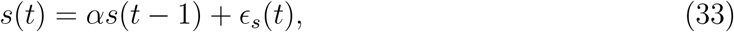

with 0 ≤ *α* < 1 and noise variable *ϵ_s_*(*t*) drawn from 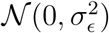. An equivalent formulation for AR(1) process is to define *α* and 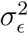 in terms of a process autocorrelation, *τ*, and process variance, 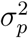, as *α* = exp(−1/ *τ*) and 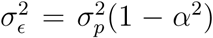. To make the process distribution stationary the correct distribution for *s*(0) is 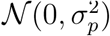. Applying this to LFADS, the prior for **u***_t_* with *t* > 1 is an independent AR(1) process in each dimension, such that for the *i^th^* element of **u***_t_* an autocorrelation *τ_i_* and process variance 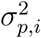 are initialized to used-defined initial values.

### 1.8 Modifications to the LFADS algorithm for stitching together data from multiple recording sessions

To accommodate multiple recordings sessions, as in **Fig. 5** of the main text, we made minor modifications to the LFADS architecture. In particular, we allowed each separate recording session to have unique input and output “adaptor” matrices. The reasons are both practical and conceptual. Practically, a different number of units are recorded in each recording session, thus the number of inputs and outputs to the LFADS algorithm needs to change accordingly. Conceptually, the hypothesis of most investigators when recording in the same area across multiple sessions is that they are recording different measurements of the same underlying (dynamical) system. Therefore, LFADS allows a different input and output transformation for each recording session to handle the different measurements, but otherwise LFADS models all the data with the same generative model, with shared parameters across all recording sessions, to allow different sessions’ measurements to improve the underlying model.

Beginning with the simpler case of the output matrices, we modified equation 5, replacing it with equation 34 to change as a function of recording session, thus introducing a session index, *s*, into the notation

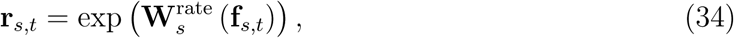

where the dimensions of 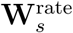 are now the number of units in the session, *D_s_*, by the number of factors in the LFADS model, *F*, which is independent of the session.

We now address the input adaptor matrices. Without multiple recording sessions, we simply fed the recorded spikes, **x***_t_* into the encoders (equations 9 and 10), single trial by single trial. To handle multiple sessions’ data, we modified this practice by introducing a per-session input adaptor matrix, 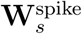. Then, for the bidirectional encoding RNN for **g**0 we modified equations 9 and 10, by inputting the linearly transformed spikes, yielding

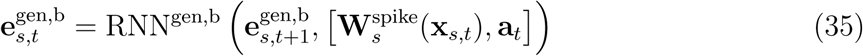

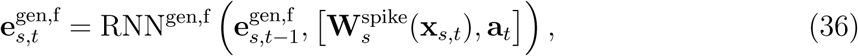

where the dimensions of the matrix is 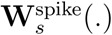 are *F* × *D_s_*. We modified the bidirectional RNN encoder for input to the RNN controller in the same way. Otherwise the LFADS architecture was identical to the standard use case, with the rest of the parameters of the LFADS architecture shared across all recording sessions.

We computed appropriate initial parameter settings for both the input adaptor and output matrices using a principal components regression technique. Briefly, we assembled a matrix of within-condition averaged firing rates for each unit across all sessions, with dimension equal to the total number of units x number of time points, 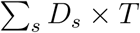. We performed principal components analysis on this matrix to reduce it to 16 principal components, equivalent to the number of factors in the model, yielding a 16 x T matrix of principal component (PC) scores. For each session, we regressed the matrix of PC scores against the condition-averaged firing rates recorded in that session. The resulting matrix of regression coefficients, which maps firing rates to PC scores, was used as the initial input adaptor matrix for that session, 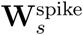. The output matrix 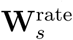 for the session was initialized to the pseudoinverse of 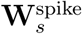. These initial adaptor and output matrices can be thought of as seeding the multi-session LFADS model with a trial-averaged correspondence across recording sessions. Note, however, that all of the output matrices and input adaptor matrices for each session were fit as parameters during training, as normal.

During training one dataset is selected at random by the algorithm (e.g. the first session), and the correct input and output adaptor matrices are then used (the matrices associated with the first session). To generate a mini-batch of gradients, the algorithm then selects a random mini-batch of data from that session and propagates it forward to evaluate the loss. The relevant gradients of the loss are then back-propagated. As a result, all shared parameters (e.g. the generator RNN parameters) are modified with every mini-batch of data regardless of dataset, while the input and output adaptor matrices are modified only when data from that session is used for training.

### 1.9 Hyper-parameters and further details of LFADS implementation

A table of the major hyper-parameters for each model is listed in **Methods Table 1**. There were a number of additional minor details that aided in the optimization and generalization of the LFADS model applied to the datasets in our study.

- To help avoid over-fitting, we added a dropout layer [14] to the inputs and to a few feed-forward (input) connections [36] in the LFADS model. Specifically, we used dropout “layers” around equation 11, around the input in equation 18, and around equation 22.
- We added an *L*_2_ penalty to recurrent portions of the generator (equations 29-32) and controller networks to encourage simple dynamics. Specifically, we regularized any matrix parameter by which **h***_t_*_−1_ was multiplied, but not those that multiplied **x***_t_*.
- As defined in eqn. 28, there is an information limiting regularizer placed on **u***_t_* by virtue of minimizing the KL divergence between the approximate posterior over **u***_t_* and the uninformative Gaussian prior.
- Following the authors in [2], we added a linearly increasing schedule on the KL divergence penalty so that the optimization does not quickly (and pathologically) set the KL divergence to 0. By 2000 steps, the schedule reached the maximum value of the KL penalty. An identical schedule was used for linearly increasing the *L*2 regularizer on the network parameters.
- We experimented with the variance of the prior distribution for the initial condition distribution and settled on a value of *κ* = 0.1, chosen to avoid saturating network nonlinearities.
- The auto-regressive prior parameters were optimized to reduce the KL divergence between inferred inputs from the approximated posterior distributions and those of the prior. In practice, nearly all AR(1) processes optimized to the uncorrelated, white noise case (*τ_i_* ≈ 0 and 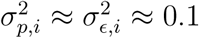). We initialized them with *τ_i_* = 10 time steps and 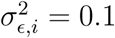.
- Unless otherwise specified, all matrices were randomly initialized with a normal distribution with mean equal to 0, and variance equal to 1*/K*, where *K* is input dimension of the matrix. All biases were initialized to 0.
- We used the ADAM optimizer, with initial learning rate of 0.01, and *β*_1_ = 0.9, *β*_2_ = 0.999, *ϵ* = 0.1. During training, the learning rate was decreased whenever the training error for the current epoch of data was greater than the last 6 training error values. In this case, the learning rate was decayed by multiplying the rate by 0.95, and 6 training epochs were required before the learning rate could be decayed again. The optimization continued until the learning rate was less than or equal to 1× 10^−5^. We routinely saved checkpoints of the model and therefore were able to capture the model with the lowest validation error.
- We clipped our hidden state **h***_t_* when any of its values went above a set threshold. This threshold was rarely hit, but was useful to avoid occasional pathological conditions.
- We used gradient clipping with a value of 200 to avoid occasional pathological gradients.
- The matrix in the **W**^fac^(·) affine transformation was row-normalized to keep the factors relatively evenly scaled with respect to each other.

**Table 1:** Important hyper-parameters of LFADS models. Listed here are the most important LFADS parameters, relating primarily to model capacity. ‘N’ - number of units in the generator, ‘F’ - number of factors, |**u***_t_*| - number of inferred inputs, ‘E’ - encoder, ‘C’ - controller, ‘G’ - generator, ‘KP’ - keep probability in dropout layers, ‘BS’ - bin size (ms).

| Model | Figure | N | F | $\vert \mathbf{u}_t\vert $ | g0 E dim | $\mathbf{u}_t$ E dim | C dim | G L2 | C L2 | KP | BS |
| --- | --- | --- | --- | --- | --- | --- | --- | --- | --- | --- | --- |
| Monkey J Maze | Main 2, 3 | 64 | 20 | 2 | 64 | 64 | 128 | 10 | 10 | 0.98 | 5 |
| Participant T5 Centerout | Main 3 | 100 | 20 | 3 | 100 | 100 | 100 | 250 | 250 | 0.98 | 5 |
| Monkey J Centerout | Main 1, 4 | 128 | 50 | 4 | 150 | 100 | 128 | 25 | 25 | 0.98 | 2 |
| Participant T7 Centerout | Main 4 | 64 | 20 | 3 | 64 | 64 | 128 | 250 | 250 | 0.95 | 5 |
| Monkey P Multi-session | Main 5 | 100 | 16 | 0 | 100 | - | - | 500 | - | 0.98 | 10 |
| Monkey P Single-session | Main 5 | 100 | 16 | 0 | 100 | - | - | 500 | - | 0.98 | 10 |
| Monkey J CursorJump | Main 6 | 128 | 50 | 4 | 150 | 100 | 128 | 25 | 25 | 0.98 | 10 |
| Lorenz attractor | Supp. 2 | 64 | 3 | 0 | 64 | - | - | 250 | - | 0.95 | a.u. |
| Chaotic RNN | Supp. 3 | 200 | 20 | 0 | 200 | - | - | 2000 | - | 0.95 | a.u. |
| Input pulses | Supp. 6, 7 | 200 | 20 | 1 | 200 | 128 | 128 | 2000 | 0 | 0.95 | a.u. |
| RNN Integrator | Supp. 8 | 200 | 20 | 1 | 128 | 128 | 128 | 2000 | 0 | 0.95 | a.u. |

**Table 2:** Signal collection technology and spike detection method.

| Model | Figure | Electrode type | Signal post-processing |
| --- | --- | --- | --- |
| Monkey J Maze | Main 2, 3 | Utah array | threshold crossing, spike sorted |
| Participant T5 Centerout | Main 3 | Utah array | threshold crossing |
| Monkey J Centerout | Main 1, 4 | Utah array | threshold crossing |
| Participant T7 Centerout | Main 4 | Utah array | threshold crossing |
| Monkey P Multi-session | Main 5 | v-probe | threshold crossing |
| Monkey P Single-session | Main 5 | v-probe | threshold crossing |
| Monkey J CursorJump | Main 6 | Utah array | threshold crossing |

### 1.10 Computing posterior averages of model variables

As the LFADS model is inherently stochastic, one needs to average in order to get good estimates of meaningful quantities within the network (e.g. the rates, **r***_t_*). For example, in de-noising a single trial of spike trains, we run the full LFADS model - both encoder and decoder on the single trial. For that single trial, we sample the stochastic variables, (eqns. 12 and 15) some number of times (e.g. 512) and then evaluate the generative portion of the model with these sampled variables. Finally, we obtain the mean of the quantity, in this case, the posterior average, computed by averaging the quantity of interest over the random samples of the stochastic variables, e.g. 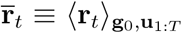. It is posterior averages such as 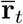 that are shown in the majority of figures.

### 1.11 LFADS related work in machine learning literature

Recurrent neural networks have been used extensively to model neuroscientific data (e.g. [31] [24], [3], [32], [28]), but the networks in these studies were all trained in a deterministic setting. An important recent development in deep learning has been the advent of the variational auto-encoder [19] [29], which combines a probabilistic framework with the power and ease of optimization of deep learning methods. VAEs have since been generalized to the recurrent setting, for example with variational recurrent networks [5], deep Kalman filters [21], and the RNN DRAW network [13].

There is also a line of research applying probabilistic sequential graphical models to neural data. Recent examples include PLDS [23], switching LDS [27], GCLDS [10], and PfLDS [9]. These models employ a linear Gaussian dynamical system state model with a generalized linear model (GLM) for the emissions distribution, typically using a Poisson process. In the case of the switching LDS, the generator includes a discrete variable that allows the model to switch between linear dynamics. GCLDS employs a generalized count distribution for the emissions distribution. Finally, in the case of PfLDS, a nonlinear feed-forward function (neural network) is inserted between the LDS and the GLM.

Gaussian process models have also been explored. GPFA [35] uses Gaussian processes (GPs) to infer a time constant with which to smooth neural data and has seen widespread use in experimental laboratories. More recently, the authors of [37] have used a variational approach (vLGP) to learn a GP that then passes through a nonlinear feed-forward function to extract the single-trial dynamics underlying neural spiking data.

Additional work applying variational auto-encoding ideas to recurrent networks can be found in [1]. The authors of [21] have defined a very general nonlinear variational sequential model, which they call the Deep Kalman Filter (DKF). The authors of [33] applied recurrent variational architectures to problems of control from raw images. Finally, [15] applied dynamical variational ideas to sequences of images. Due to the generality of the equations in many of these references, LFADS is likely one of many possible instantiations of a variational recurrent network applied to neural data (in the same sense that a convolutional network architecture applied to images is also a feed-forward network, for example).

The LFADS model decomposes the latent code into an initial condition and a set of innovation-like inferred inputs that are then combined via an RNN to generate dynamics that explain the observed data. Recasting our work in the language of Kalman filters, our nonlinear generator is analogous to the linear state estimator in a Kalman filter, and we can loosely think of the inferred inputs in LFADS as innovations in the Kalman filter language. However, an “LFADS innovation” is not strictly defined as an error between the measurement and the readout of the state estimate. Rather, the LFADS innovation may depend on the observed data and the generation process in extremely complex ways.

## 2 Synthetic datasets

### 2.1 Lorenz system

The Lorenz system is a set of nonlinear equations for three dynamic variables. Its limited dimensionality allows its entire state space to be visualized. The evolution of the system’s state is governed as follows

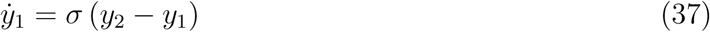

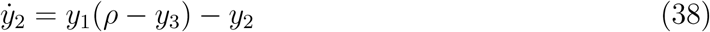

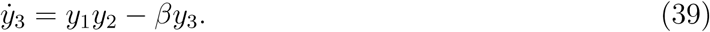

We used the standard parameter values known for inducing chaos, *σ* = 10, *ρ* = 28, and *β* = 8/3, and used Euler integration with Δ*t* = 0.006. As in [37], we simulated a population of neurons with firing rates given by linear readouts of the Lorenz variables using random weights, followed by an exponential nonlinearity. Spikes from these firing rates were then generated by a Poisson process.

Our synthetic dataset consisted of 65 conditions, with 20 trials per condition. Each condition was obtained by starting the Lorenz system with a random initial state vector and running it for 1s. Twenty different spike trains were then generated from the firing rates for each condition. Models were trained using 80% of the data (16 trials/condition) and evaluated using 20% of the data (4 trials/condition). While this simulation is structurally quite similar to the Lorenz system used in [37], we purposefully chose parameters that made the dataset more challenging. Specifically, relative to [37], we limited the number of observations to 30 simulated neurons instead of 50, decreased the baseline firing rate from 15 spikes/sec to 5 spikes/sec, and sped up the dynamics by a factor of 4.

### 2.2 Chaotic RNNs as data generators

We tested the performance of each method at inferring the dynamics of a more complex nonlinear dynamical system, a fully recurrent nonlinear neural network with strong coupling between the units. We generated a synthetic dataset from an *N*-dimensional continuous time nonlinear, so-called, “vanilla” RNN,

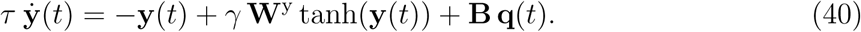

This makes a compelling synthetic case study for our method because many recent studies of neuroscientific data have used vanilla RNNs as their modeling tool (e.g. [31] [24], [3], [32], [28]). It should be stressed that the vanilla RNN used as the data RNN here does not have the same functional form as the network generator used in the LFADS framework, which is a GRU (see section 1.6). For experiments in **Supp. Fig. 3**, we set **B** = **q** = 0, but we included an input for experiments in **Supp. Fig. 6**.

The elements of the matrix **W**^y^ were drawn independently from a normal distribution with zero mean and variance 1/*N*. We set *γ* to either 1.5 or 2.5, both of which produce chaotic dynamics at a relatively slow timescale compared to *τ* (see [31] for more details). The smaller *γ* value produces “gentler” chaotic activity in the data RNN than the larger value. Specifically, we set *N* = 50, *τ* = 0.025 s and used Euler integration with Δ*t* = 0.01 s. Spikes were generated by a Poisson process with firing rates obtained by scaling each element of tanh(**y**(*t*)) to take values in [0, 1], and then used as the rate in a Poisson process to give rates lying between 0 and 30 spikes/s.

Our dataset consisted of 400 conditions obtained by starting the data RNN at different initial states with elements drawn from a normal distribution with zero mean and unit variance. Firing rates were then generated by running the data RNN for 1 s, and 10 spiking trials were produced for each condition, yielding a total of 4,000 spiking trials. Models were trained using 80% of the data (8 trials/condition) and evaluated using 20% of the data (2 trials/condition).

### 2.3 Inferring pulse inputs to a chaotic RNN

We tested the ability of LFADS to infer the input to a chaotic RNN (**Supp. Figs. 6,7**). In general, the problem of disambiguating dynamics from inputs is ill-posed, so we encouraged the dynamics to be as simple as possible by including an L2 regularizer in the LFADS network generator (see **Methods Table 1**). We note that weight regularization is a standard technique that is nearly universally applied to neural network architectures.

Focusing on **Supp. Fig 6**, we studied the synthetic example of inferring the timing of a delta pulse input to a randomly initialized RNN. To introduce an input into the data RNN, the elements of **B** were drawn independently from a normal distribution with zero mean and unit variance. During each trial, we perturbed the network by delivering a delta pulse of magnitude 50, *q*(*t*) = 50*δ*(*t* − *t*_pulse_), at a random time *t*_pulse_ between 0.25s and 0.75s (the full trial length was 1s). This pulse affects the underlying rates produced by the data RNN, which modulates the spike generation process. To test the ability of the LFADS model to infer the timing of these input pulses, we included in the LFADS model an inferred input with dimensionality of 1. We explored the same two values of *γ* as in the synthetic example to model chaotic RNN dynamics, 1.5 and 2.5. Other than adding the input pulses, the data for input-pulse perturbations were generated as in the first data RNN example described above.

After training, which successfully inferred the firing rates, we extracted inferred inputs from the LFADS model (eqn. 15) by running the system 512 times for each trial, and averaging, defining 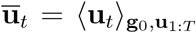. To see how the timing of the inferred input was related to the timing of the actual input pulse, we determined the time at which 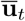 reached its maximum value.

### 2.4 Inferring white noise input in an RNN trained to integrate to bound

We tested the ability of LFADS to infer the input to a vanilla RNN trained to integrate a noisy signal to a +1 or −1 bound. The signal was drawn from a Gaussian distribution with zero mean and variance 0.0625. Weight matrices for the data RNN were drawn independently from a Gaussian distribution with zero mean and variance 0.64/*N*, and L2 regularization was used during training. 800 conditions were generated with white noise inputs, and 5 spiking trials were generated per condition. This resulted in 4,000 1s spiking trials. 3,200 trials were used for training and 800 trials were used for validation.

After training LFADS on the integrate to bound data, inferred inputs (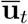) were extracted by averaging over 1024 runs of each trial. These inferred inputs were then compared (using *R*^2^) with the real inputs to the integrate to bound model, which were saved down previously during training.

## 3 Neural datasets - Research participants with paralysis

Permission for these studies was granted by the US Food and Drug Administration (Investigational Device Exemption) and Institutional Review Boards of Stanford University (protocol # 20804), Partners Healthcare/Massachusetts General Hospital (2011P001036), Providence VA Medical Center (2011–009), and Brown University (0809992560). The participants in this study were enrolled in a pilot clinical trial of the BrainGate Neural Interface System (http://www.clinicaltrials.gov/ct2/show/NCT00912041). Informed consent, including consent to publish, was obtained from the participants prior to their enrollment in the study.

Participant T7 was a right-handed man, 54 years old at the time of the research sessions reported here, who was diagnosed with ALS and had resultant motor impairment (ALSFRS-R of 17). In July 2013, participant T7 had two 96-channel intracortical silicon micro-electrode arrays (1.5 mm electrode length, Blackrock Microsystems, Salt Lake City, UT) implanted in the hand area of dominant motor cortex. T7 retained very limited and inconsistent finger movements. Data reported are from T7’s post-implant day 231.

A second study participant, T5, is a right-handed man, 63 years old at the time of the research sessions reported here, with a C4 ASIA C spinal cord injury that occurred approximately 9 years prior to study enrollment. He retains the ability to weakly flex his left (non-dominant) elbow and fingers; these are his only reproducible movements of his extremities. He also retains some slight residual movement which is inconsistently present in both the upper and lower extremities, mainly seen at ankle dorsiflexion and plantarflexion, wrist, fingers and elbow, more consistently present on the left than on the right. Occasionally, the initial slight voluntary movement triggers involuntary spastic flexion of the limb. In Aug. 2016, participant T5 had two 96-channel intracortical silicon micro-electrode arrays (1.5 mm electrode length, Blackrock Microsystems, Salt Lake City, UT) implanted in the upper extremity area of dominant motor cortex. Data reported are from T5’s post-implant day 51.

### 3.1 Task design and data analysis

Neural data were recorded during “Center-out-and-back” target acquisition tasks. The data were originally collected for neural prosthetic decoder calibration, as part of research testing algorithms for closed-loop neural cursor control ([12], [25], [26]). In the Center-out-and-back task, data were collected either in motor-based control (with T7, who retained limited residual movements), or an attempted movement paradigm (with T5, who did not retain sufficient movement to reliably measure or physically control a cursor). In motor-based control, T7 controlled the position of a cursor on a computer screen by making physical movements with his fingers on a wireless touch-pad (Magic Trackpad; Apple, Cupertino, CA). The cursor began in the center of the screen, and targets would appear in one of 8 locations on the periphery. The participant then acquired the targets by moving the cursor over the target and holding it over the target for 500 ms. Participant T7’s limited movements spanned a small region on the touch-pad, approximately 1/8”–1/4” wide. In the attempted movement paradigm, the cursor was automatically moved directly toward the target by the computer, and T5 was asked to attempt movements of his whole arm that followed the movements of the cursor.

Voltage signals from each of the electrodes were band-pass filtered from 250 to 7500 Hz and then processed to obtain multi-unit ‘threshold crossings,’ i.e., discrete events that occurred whenever the voltage crossed below a threshold (choice of threshold was dependent on the array-T7 lateral array: -80 *μ*V; T7 medial array: -95 *μ*V; T5, both arrays: -3.5 times the r.m.s. voltage on each channel.). For the present analyses, we did not “spike sort” and instead grouped together threshold crossings on a given electrode. These spikes therefore can include both single- and multi-unit activity. For both participants, analysis was restricted to channels known to show significant modulation during movement attempts (T7: 78 channels; T5: 187 channels).

Neural control and task cueing were controlled by custom software run on the Simulink/xPC real-time platform (The Mathworks, Natick, MA), enabling millisecond-timing precision for all computations. Neural data collected by the NeuroPort System (Blackrock Microsystems, Salt Lake City) were available to the real-time system with 5-ms latency. Visual presentation was provided by a computer via a custom low-latency network software interface to Psychophysics Toolbox for MatLab and an LCD monitor with a refresh rate of 120 Hz. Frame updates from the real-time system occurred on screen with a latency of approximately 7 ± 5 ms.

## 4 Neural datasets - Nonhuman primates

All procedures and experiments were approved by the Stanford University Institutional Animal Care and Use Committee.

### 4.1 Maze task

An adult male macaque monkey (monkey J) was trained to sit head-fixed in a primate chair and perform 2D target acquisition tasks in a fronto-parallel plane by controlling an on-screen cursor with his hand movements. Monkey J was implanted with two 96-electrode arrays (1 mm electrodes spaced 400 *μm* apart, Blackrock Microsystems) using standard neurosurgical techniques. The arrays were implanted into M1 and dorsal premotor cortex (PMd) of the hemisphere contralateral to his reaching arm.

The Maze task is a variant of a center-out delayed reach task, whose details have previously been described [17]. Briefly, monkey J made arm movements in a 2-dimensional workspace while the position of the right index and middle fingertips was tracked optically. This tracked position controlled the movements of a virtual cursor, and the cursor’s position floated 2.5 cm above the hand. To initiate a trial, the monkey fixated on a fixation spot for >400 ms, after which a target appeared. After a delay period (varying from 0 - 900 ms), a go cue instructed the monkey to begin his movement. A set of virtual barriers in the workspace facilitated the instruction of curved or straight reach trajectories. Contact with a barrier resulted in an unrewarded trial. A trial was counted as a success, and reward delivered, if the monkey held the cursor on the target for 450 ms.

Several de-noising methods were applied to the Maze dataset. For all methods, individual trials were aligned to movement onset (the point at which movement is first detectable), and data consisted of 450 ms preceding and following movement onset (for a total of 900 ms per trial). The dataset consisted of 2296 trials across 108 different reach conditions (target and barrier locations), and 202 single units were isolated from the recorded activity.

### 4.2 Center-out and Cursor Jump tasks

These experiments were also performed with Monkey J. Experiments were controlled using custom MatLab and Simulink Realtime software (Mathworks, USA). Arm reaches were made with the display blocking the monkey’s view of his hand. The task was displayed in virtual reality using a Wheatstone stereograph with a latency of 7 ± 4 ms as described in ([11]). The virtual computer cursor followed the velocity of a reflective bead taped to the monkey’s hand, which was tracked via an infrared system at 60 Hz (Polaris, Northern Digital, Canada). The non-reaching arm was gently restrained. To successfully acquire a target, the monkey had to hold the cursor within a 4 x 4 cm target acquisition area for a continuous 500 ms. A target color change cued that the cursor was within the acquisition area. If the cursor left the target area during this hold period, the 500 ms timer reset. The monkey had to acquire the target within a time limit of 2 seconds to receive a liquid reward and success tone.

Voltage signals from each of the electrodes were band-pass filtered from 250 to 7500 Hz and then processed to obtain multi-unit ‘threshold crossings’, i.e. discrete events that occurred whenever the voltage crossed below a threshold (set at the beginning of each day to be -4.5 times r.m.s. voltage). For the “Center-out-and-back” and “Cursor Jump” tasks, we did not spike sort the data and instead grouped together threshold crossings on a given electrode. These threshold crossing events therefore can include both single- and multi-unit activity.

For the LFP (**Fig. 4** of main text) and Cursor Jump analyses (**Fig. 6** of main text), data analyzed were from dataset 2015-04-15, which occurred 69 months after the implantation of recording arrays. A single LFADS model was fit to data from two types of reaching tasks – a standard “Center-out-and-back” task and a Cursor Jump task.

In the Center-out-and-back task, targets alternated between being located at the workspace center or at a randomly chosen target out of 8 possible target locations, all 12 cm away from the workspace center and evenly spaced around a circle. In the Cursor Jump task, targets were located either at the workspace center or one of two radial target locations located 12 cm away from the workspace center, in opposite directions. The three possible targets lay along the vertical monitor axis.

The ‘cursor jump’ manipulation at the heart of the Cursor Jump Task was applied on a random 25% of trials towards radial targets. On these randomly selected perturbation trials, the cursor position was offset by 6 cm perpendicular to the vertical axis. The jump happened after the cursor traveled 6 cm towards the target along the vertical axis. Only one perturbation occurred per trial. The time when the cursor jump command was sent to the display computer was recorded with 1 ms resolution, after which it appeared at the next 120 Hz monitor update. The delivery of cursor jump position offsets required us to counteract this offset at the end of each perturbed outward trial so as to not carry a (possibly accumulating) hand-to-cursor offset over multiple trials. Thus, we applied a second, opposite cursor jump as soon as the center target re-appeared, resulting in a consistent hand-to-cursor position relationship at the start of each outward trial.

To train the LFADS model, spike trains were binned at 10 ms resolution. A single LFADS model was fit to a combined dataset containing center-out-and-back trials (8 targets), outward trials without perturbations (2 targets), outward trials with perturbations (2 targets, 2 perturbation directions), and return-to-center trials from the perturbed/unperturbed outward trials, for a total of 5140 trials. 800 ms of data were taken for each trial, with data aligned to the start of the trial (target onset). In cases of perturbations, most jumps happened between 400-550 ms post-target onset. The model was allowed to infer 4 inputs to the generator in order to fit the data.

### 4.3 Multi-session V-probe recordings

One adult male macaque monkey (P) was trained in a behavioral task as described below. After initial training, we performed a sterile surgery during which the macaque was implanted with a head restraint and a recording cylinder (NAN Instruments), which was located over left, caudal, dorsal premotor cortex (PMd). The cylinder was placed surface normal to the skull and secured with methyl methacrylate. A thin layer of methyl was also deposited atop the intact, exposed skull within the chamber. Before recording sessions began, a miniature craniotomy (3 mm diameter) was made under ketamine/xylazine anesthesia, targeting an area in PMd which responded during movements and palpation of the upper arm (17 mm anterior to interaural stereotaxic zero).

In the behavioral task, monkey P was trained to use his right hand to grasp and translate a custom 3D printed handle (Shapeways, Inc.) attached to a haptic feedback device (Delta.3, Force Dimension, Inc.). The other arm was comfortably restrained at the monkey’s side. The haptic device was controlled via a 4 kHz poll position, update force feedback loop implemented in custom software written in C++ atop Chai3D (http://chai3d.org). The weight of the device was compensated by upward force precisely applied by the device’s motors, such that the motion of the device felt nearly effortless because the device’s mechanical components were lightweight and had low inertia. The device endpoint with the attached monkey handle was constrained via software control to translate freely in the fronto-parallel plane. The handle was custom 3D printed and contained a beam break detector which indicated whether the monkey was gripping the handle. The task was controlled using custom code running on a dedicated Simulink Real Time operating system. Hand position was recorded at 1 kHz, and the 2D position of the device was used to update the position of a white circular cursor at the refresh rate of 144 Hz with a latency of 4-12 ms (verified via photodiode) displayed on an LCD screen located in front of the monkey and above the haptic device, in the same fronto-parallel plane as the device itself. The display was driven by custom software driven by Psychophysics Toolbox. A plastic visor was used to mask the monkey’s visual field such that he could see the screen but not his hand or the haptic device handle.

The monkey was trained to perform a delayed center-out reaching task by moving the haptic device cursor towards visual targets displayed on the screen. Monkeys initiated the task by holding onto the device handle, which was detected by a beam break photodiode built into the handle. At the start of each trial, the device gently returned the hand to the center position and supported the weight of the arm from below in that position (by rendering a narrow virtual shelf just below the haptic cursor). At *target onset*, one or more reach targets appeared as hollow circles at one of 8 radial locations located 10 cm from the position. After a variable delay period (50-800 ms), the *go cue* was indicated visually by the target outline filling in with color. A trial was successful if the monkey remained still during the delay period, initiated the reach within 600 ms after the go cue, and held in the reach target for 50 ms. In some sessions, the monkey performed additional trial conditions with different target locations or forces applied to the haptic device. These trials were excluded from analysis; only successful center-out reaches were included. Hand velocities were computed by applying a smoothing, differentiating filter (Savitzy-Golay, 2nd-order, 3 ms smoothing widow) to the raw position time series. Reaction time was measured from the visual display of the go cue detected at the photodiode until the hand speed in the fronto-parallel plane reached 5% of the peak speed on each trial.

Electrophysiological recordings were performed by slowly lowering a linear multielectrode array with 24 recording channels (Plexon V-probe or U-probe) to a position where the channels likely spanned the layers of the cortex based on properties of the neural signals. We allowed 45-90 minutes to allow the probe to settle before beginning experiments. All 24 channels were amplified and sampled at 30 kHz (Blackrock Microsystems), high-pass filtered (fourth-order Butterworth filter, 250 Hz corner frequency), and thresholded at -3.5x RMS voltage on each channel. Threshold crossings on adjacent channels that occurred within 0.5 ms of each other were removed from one of the channels to avoid duplicate detection of spiking along the array. Threshold crossing rates were then binned at 10 ms on each channel.

Experimental sessions were screened on the basis of minimum trial count (200 trials); one dataset was manually excluded on the basis of an abrupt discontinuity in the recorded firing rates over the session. Following this screening, a total of 44 consecutive experimental sessions were included, comprising recording locations in the upper arm representation of primary motor cortex and dorsal premotor cortex. A 1200 ms time window beginning 500 ms before the go cue to 700 ms afterwards was chosen from each successful trial and used to train the LFADS model.

## 5 Analysis Methods by Figure

We used a number of analysis methods on either smoothed neural data, or the output of LFADS, typically the rates, factors or inferred inputs. All of these analyses methods are standard, but we provide references and operating parameters here.

### 5.1 Fig. 2 - Kinematic predictions on the Maze dataset

We used Optimal Linear Estimation [30] to create decoders that mapped neural features onto the measured x and y reaching velocities. The inputs to the decoder were the raw or de-noised neural data from 250 ms prior to 450 ms post movement onset. De-noising was achieved via one of three techniques: Gaussian smoothing, GPFA[35], or LFADS. For Gaussian smoothing, the millisecond-binned spike trains were convolved with a Gaussian function with standard deviation (s.d.) of 40 ms. For GPFA and LFADS, millisecond-binned spike trains were re-binned at 5 ms resolution, and both techniques were allowed to fit 20 latent dimensions (factors). The neural features from each technique were the Gaussian-smoothed firing rates, factor estimates using GPFA, or de-noised firing rates using LFADS. In all cases, to decode kinematics, the neural features were ‘lagged’ by 90 ms to account for delays between neural activity and measured kinematics, and the neural features were binned at 20 ms (the parameters of Gaussian s.d., and lag were optimized using cross-validated decoding).

Kinematic predictions were generated using 5-fold cross-validation. The subsampling analyses followed the above, with limited populations achieved via random subsampling (without replacement) from the full population of 202 neurons. Decoding performance was quantified using goodness of fit (*R*^2^) between the original and reconstructed velocities (validation trials from the 5-fold cross-validated decoding) for the x and y dimensions. For the sample reconstructed reach trajectories shown in Fig. 2a, trajectories were seeded with the true initial position, and subsequent points in the trajectory were calculated by simply integrating the decoded velocity at each timestep.

Note that for offline decoding analyses, the approach of smoothing neural data and then linearly regressing against kinematics, outlined here, is a generalization of common brain-machine interface (BMI) decoding approaches such as the Kalman Filter. This relationship is outlined in [34]; briefly, the Kalman filter can be viewed as a two step process - first smoothing the data, and subsequently performing a linear dimensionality reduction step that maps the smoothed, high-dimensional neural data onto kinematics. In the Kalman Filter the amount of smoothing is largely determined by the simple linear dynamical system (LDS) that models state evolution (i.e., models changes in kinematics). This can be especially problematic in datasets with highly varied kinematics, such as the complex “maze” reaching dataset, where a simple LDS does not provide a good model of observed kinematics. Therefore, to avoid having the degree of smoothing influenced by a poorly-fit kinematics model, we optimized the smoothing parameter using cross-validated decoding as described above.

Further improvement can be achieved for online (closed-loop) BMI control using an additional “intention estimation” step, and then regressing neural data against the inferred intention rather than the measured kinematics. This “intention estimation” step has been shown to improve closed-loop BMI control when intention is estimated from hand reaching data (e.g. the FIT Kalman Filter, [7]) or estimated from closed-loop BMI control (e.g. the Re-FIT Kalman Filter, [11, 12]). However, to date, these approaches having been applied to simple datasets (point-to-point movements) in order to calibrate a BMI decoder, and make the assumption that the subject’s intention was to move in a straight line toward the target. In the complex “maze” dataset analyzed in Fig. 2, the monkey made curved reaches which violate this assumption - therefore our decoding approach used regression against measured kinematics rather than attempting to infer the subject’s intention.

### 5.2 Fig. 3 - Rotations in state space

Rotations in state space were found using the jPCA technique, whose mathematical details are presented elsewhere [6]. We briefly summarize the overall approach here. jPCA was applied in two ways: first to examine rotations in the condition-averaged responses, and subsequently for the single trial responses. For condition-averaged responses, each neuron’s response was first averaged across all trials of the identical condition to create a set of condition-averaged firing rates. These firing rates were smoothed via convolution with a Gaussian kernel, with the width of the kernel chosen to reduce the noise in the firing rates without smoothing away the rotational content. Smoothed firing rates were then mean-centered across conditions at every time point by subtracting the average across-condition response from the response of each individual condition. The mean-centered rates were then “soft-normalized” [6] to prevent individual neurons (e.g. high firing rate or potentially noisy neurons) from dominating the results of the subsequent dimensionality-reduction step. These high-dimensional neural firing rates were projected into a low-dimensional subspace using PCA. Within this subspace (neural state space), we then used the jPCA technique to find planes that are best fit by a linear dynamical system with purely rotatory dynamics.

For the subsequent single trial responses, the goal was to examine the same rotations in state space that were found via condition averaging, but examine their consistency at the level of single trials. Therefore, the single trial data was projected into jPCA planes via the projections that were calculated in the condition averaged analysis.

For monkey J, all trials were aligned to movement onset. We used 250 ms for jPCA analysis, with the time window starting 60 ms prior to movement onset. Observed neural firing rates were smoothed with a 40 ms s.d. Gaussian kernel to reduce noise, and soft-normalized with a value of 0.1. For the de-noised LFADS data, further smoothing and de-noising had little effect, so the parameters used were a 25 ms s.d. Gaussian kernel with a negligible soft-normalization value (5e-5). For the initial dimensionality-reduction step (PCA), 10 PCs were kept and used for jPCA.

As with the monkeys, the rotations in state space for research participants with paralysis are found by identifying the time period starting just before the rapid change in neural activity that occurs with a movement attempt [25]. For participant T5, because no movement was measurable, data were simply aligned to the start of the trial (i.e., the point at which targets are displayed). The window taken for jPCA analysis was 400 ms of data beginning 240 ms after the start of the trial. As with the monkey data, larger parameters for smoothing and greater soft-normalization were used to de-noise the observed neural responses, vs. the LFADS de-noised neural responses. These were a Gaussian kernel s.d. and soft-normalization parameter of 40 ms and 10 for the observed responses, and 25 ms and 5 for the LFADS de-noised neural responses.

### 5.3 Fig. 4. - LFP analysis

For both human (participant T7) and monkey (J) data, recorded LFP was originally sampled with high bandwidth (human: 30 kHz, monkey: 2kHz). Human data was digitally re-referenced using common-average referencing to remove global noise artifacts. Human and monkey data were low-pass filtered with a 75 Hz cutoff frequency using a 4th order Butterworth filter to minimize the contribution of action potentials to the LFP signal. Both a forwards and backwards pass of the filter (i.e., acausal filtering) were used in order to minimize group delay. Data were then filtered again with an anti-aliasing filter (8th order Chebyshev Type I lowpass filter with cutoff of 0.8 * sampling frequency / 2) and then resampled to 1 kHz for all subsequent analyses. Data analyzed were from a center-out-and-back movement paradigm. Participant T7 made movements of his index finger on a touchpad to control a cursor’s on-screen movements. Monkey J made movements of his hand in free space to control the movements of a cursor. Data analyzed were from the first 300 ms (participant T7) or 250 ms (monkey J) after target onset. For each recording channel on the electrode arrays, cross-correlograms were computed between the measured spiking activity and the recorded local field potentials on the same electrode, on a single trial basis. Cross-correlograms were then averaged across all trials. For the shuffle analyses, spiking data from an individual trial was cross-correlated with LFP data from a random trial, and these correlograms were averaged across trials.

### 5.4 Fig. 5 - Kinematic predictions of LFADS multi-session and single-session models

We used optimal linear estimation to create decoders to predict x and y reaching velocities. For decoding from LFADS, we used the factors rather than the predicted firing rates, as the neurons recorded on an individual session could unevenly represent the full set of reaching directions well, even if the underlying factors from which the rates are extracted represent all directions evenly. Aside from using factors rather than rates, the inputs to the decoder were prepared and the performance evaluated as described in 5.1. For single-dataset LFADS models, we fit individual decoders to map from each model’s factors to x and y velocities. For the stitched multi-session LFADS model, a single decoder was fit and cross-validated on all datasets simultaneously. We then computed the goodness of fit (*R*^2^) and averaged across x and y velocities.

For reaction time prediction, we used a largely unsupervised method previously described in [18]. Briefly, for each of the single-session models and the multi-session model, we performed demixing principal components analysis [20] on the factor outputs. We then projected the factors along the highest-variance, condition-independent mode, and normalized the projection to a range of 0 to 1. We then took the time at which this projection crossed a certain threshold on each trial to be the predicted reaction time, and computed the correlation coefficient between predicted and actual reaction times. For each model, we then optimized only the threshold to maximize the correlation coefficient between time of threshold crossing and reaction time.

### 5.5 Fig 6 - tSNE visualization for CursorJump data

The pattern of inputs inferred by LFADS for individual trials were mapped into a 2-dimensional space using t-Distributed Stochastic Neighborhood Embedding (t-SNE [22]). Data were aligned to the time of perturbation for perturbed trials or the mean perturbation time for the given target direction for unperturbed trials (407 ms for downward targets, 487 ms for upward targets). t-SNE mapped using the t-SNE toolbox for MatLab (https://lvdmaaten.github.io/tsne/). Inferred inputs were calculated via posterior averaging, as described in section 1.10. LFADS inferred the input values at 10 ms resolution (i.e., the resolution at which the neural data was binned before being passed into LFADS). These values were then smoothed using a causal Gaussian filter with a 20 ms standard deviation. Data fed into t-SNE consisted of the inferred input values from 40 ms to 240 ms after the time at which the task perturbation occurred (or after the mean perturbation time for unperturbed trials, as described above). t-SNE initially pre-processes data by reducing its dimensionality via PCA, and the dimensionality of the pre-processed data was chosen to be 30 dimensions. The t-SNE perplexity parameter was set to 30, and sweeping this parameter between 10 to 50 had little qualitative effect on the discernibility of the three data clusters.

## Supplemental Figures

**Figure 1:**
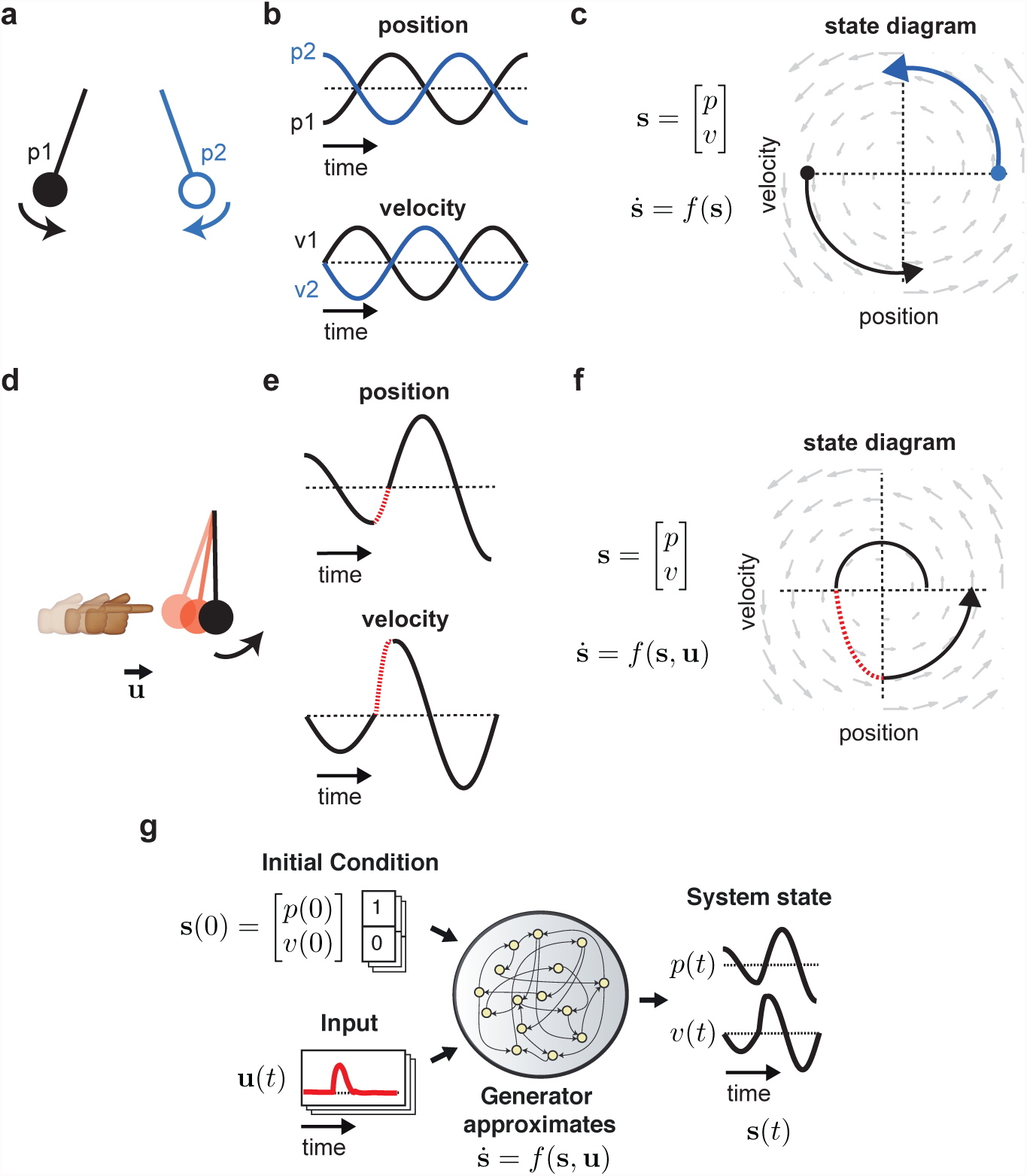
Conceptual dynamical system: 1-D pendulum. **a** A 1-dimensional pendulum released from position *p*_1_ or *p*_2_ has different positions and velocities over time. **b** The state of the pendulum is captured by a 2-D vector that specifies its position and velocity, shown here evolving as a function of time (blue and black traces correspond to different initial conditions *p*_1_ and *p*_2_). **c** The evolution of the state follows dynamical rules, i.e. the pendulum’s equations of motion. In this autonomous dynamical system, i.e. there are no perturbations, knowing the pendulum’s initial state (filled circles) and dynamical rules that govern the state evolution (gray vector field) gives full knowledge of the pendulum’s state as a function of time (black and blue traces). **d-f** Perturbations to the pendulum, an example of input-driven dynamics. **d** The pendulum’s motion might be perturbed by an external force, e.g. it is bumped rightward. **e,f** With a perturbation, the evolution of the system’s state during the perturbation no longer follows its autonomous dynamical rules. This is shown as dashed red lines in the position vs. time and velocity vs. time plots, as well as the state-space diagram. A perturbation can be modeled by transforming the equations to allow an input term **u**(*t*) that models the perturbation. **g** LFADS modeling of the perturbed pendulum. Traces of the pendulum motion are used to train LFADS. During training LFADS’ generator learns to approximate the pendulum dynamics, 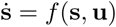, using its own internal state and dynamics. LFADS learns a per-trial initial generator state **g**_0_, which allows it to model trials that start with different initial pendulum states, **s**_0_. LFADS also learns a set of time-varying inputs per-trial, which allows it to model perturbations to the pendulum system, **u**(*t*). These three pieces of information are enough to reconstruct each trial.

**Figure 2:**
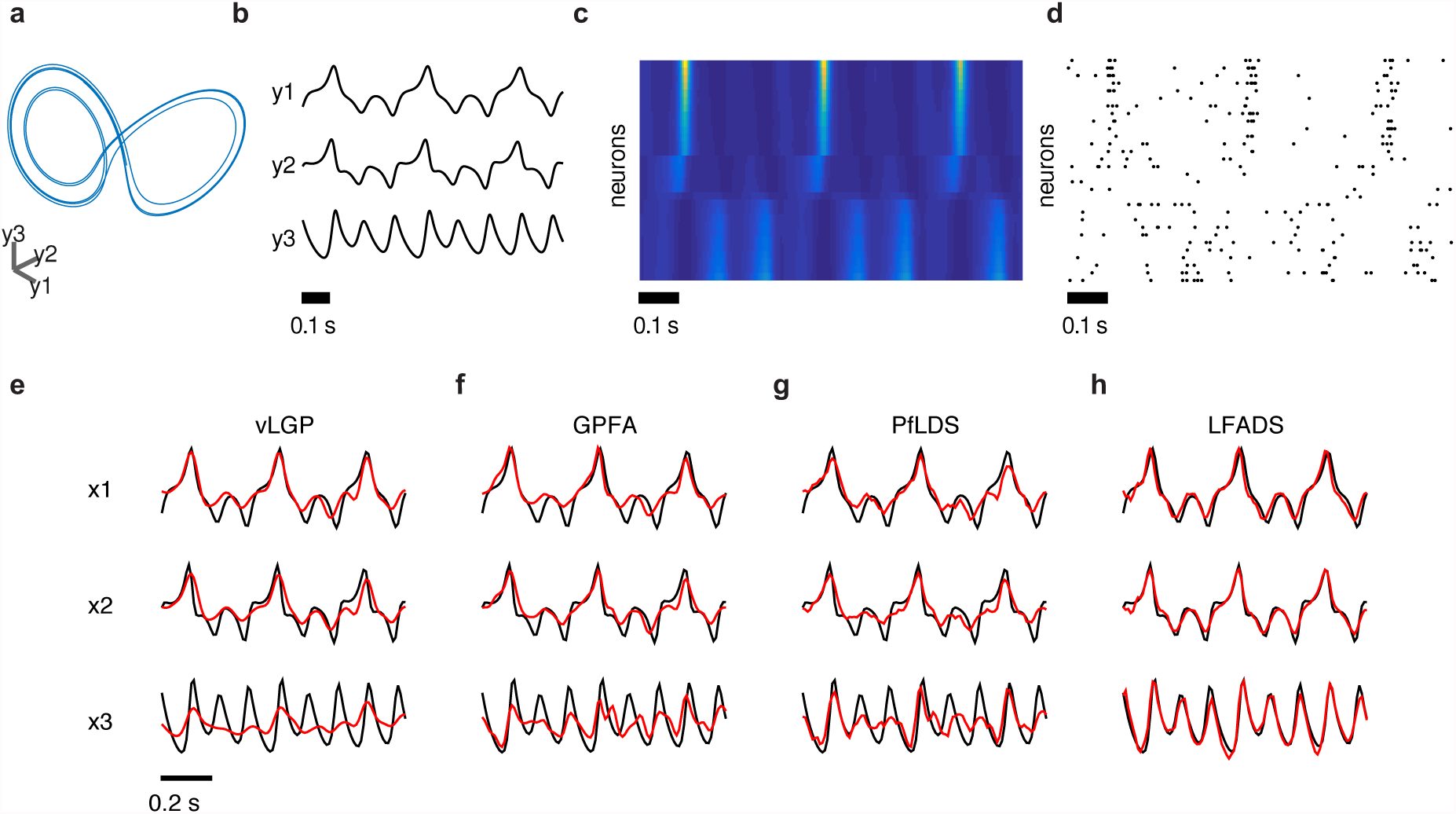
LFADS applied to Lorenz attractor. We compared the performance of LFADS to three existing methods that estimate latent state from neural data: Variational Latent Gaussian Process (vLGP[7]), Gaussian Process Factor Analysis (GPFA[6]), and Poisson Feed-forward neural network Linear Dynamical System (PfLDS[1]). To test LFADS and to compare its performance with other approaches, we generated synthetic stochastic spike trains from deterministic nonlinear systems. The first is the standard Lorenz system (see Online Methods for equations and details). **a** An example trial illustrating the evolution of the Lorenz system in its 3-dimensional state space and **b** its dynamic variables as a function of time. **c** Firing rates for the 30 simulated neurons are generated by a linear readout of the latent variables followed by an exponential nonlinearity, with neurons sorted according to their weighting for the first Lorenz dimension. **d** Spike times for the neurons are generated from the rates of the simulated neurons. **e-h** Sample performance for each method applied to spike trains based on Lorenz attractor. Each panel shows actual (black) and inferred (red) values of the three latent variables for a single example trial for the 4 methods: **e** vLGP, **f** GPFA, **g** PfLDS, **h** LFADS. For LFADS, posterior means were averaged over 128 samples of **g**_0_ conditioned on the particular input sequence.

**Table 1:** Performance of various methods on Lorenz attractor. vLGP [7], GPFA [6], and PfLDS [1]. LFADS recovers more variance of the latent Lorenz dynamics, as measured by *R*^2^ between the linearly transformed output of each model, and the dynamics of the latent Lorenz dimensions. We quantify this using *R*^2^, i.e., the fraction of the variance of the actual latent variables captured by the estimated latent values. For each method, the inferred latents of the 30 simulated neurons were linearly transformed to the actual 3D Lorenz latents to facilitate direct comparison. As shown, LFADS accurately recovered the latent dynamics underlying the observed spike trains, consistently outperforming the three other methods.

| Dim | vLGP | GPFA | PfLDS | LFADS |
| --- | --- | --- | --- | --- |
| y1 | 0.761 | 0.713 | 0.784 | **0.840** |
| y2 | 0.688 | 0.725 | 0.732 | **0.921** |
| y3 | 0.218 | 0.325 | 0.368 | **0.872** |

**Figure 3:**
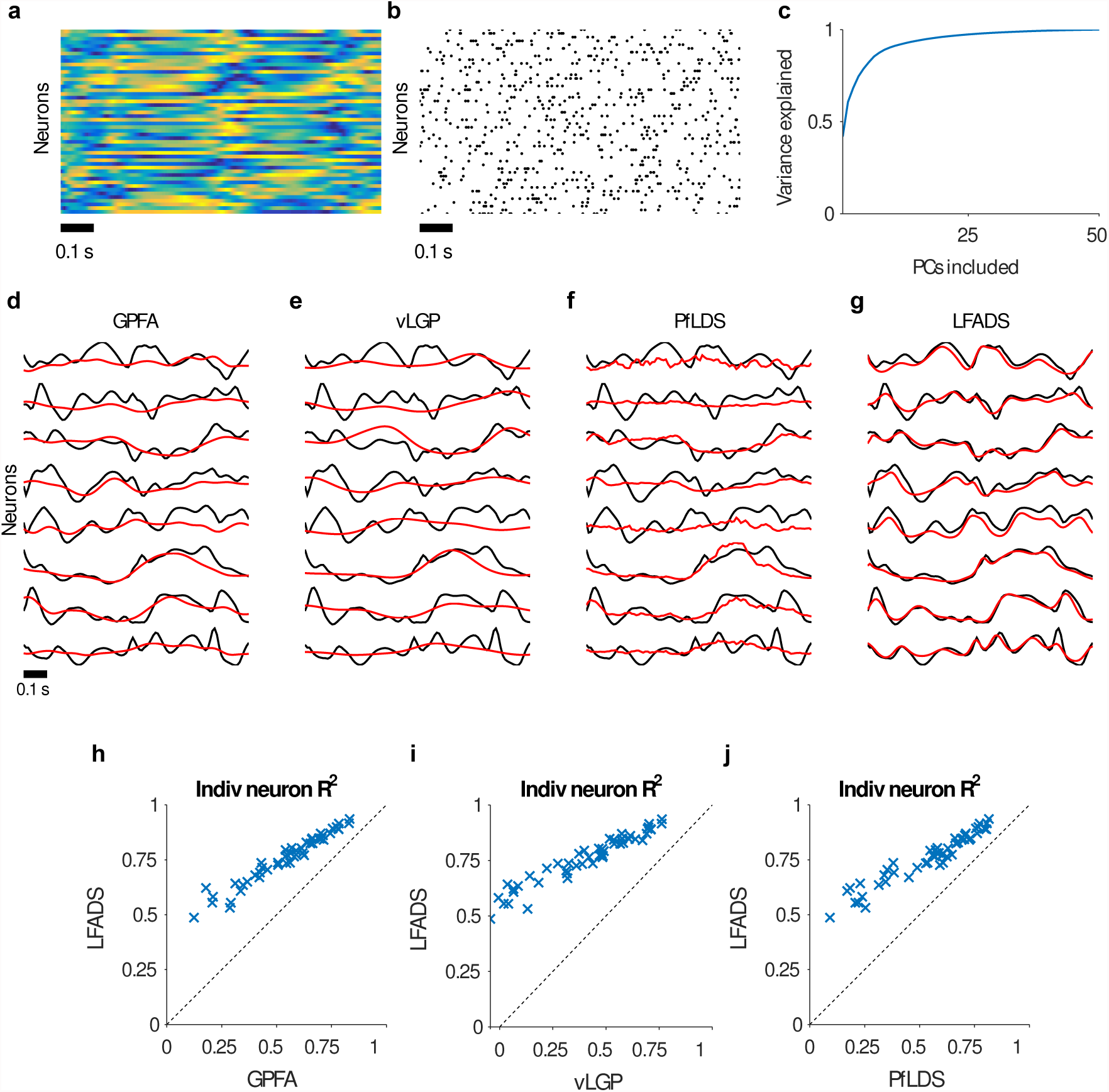
LFADS applied to autonomous chaotic data RNN. **a-c** We generated high-dimensional chaotic dynamics from an RNN. **a** Firing rates generated on one example trial by the chaotic data RNN (colors show rates fluctuating between -1 and 1). **b** The resulting spikes emitted from a Poisson process whose underlying rates were the normalized continuous rates of the RNN. **c** We used principal components analysis to assess the dimensionality of the data. As expected, the state of the data RNN had lower dimension than its number of neurons, and 20 principal components were sufficient to capture > 95% of the variance of the system. So we restricted the latent space to 20 dimensions for each of the models tested and, in the case of LFADS, set the dimensionality of temporal factors to 20 as well (*F* = 20). **d-g** Sample performance for each method on the RNN task. We tested the performance of the methods at extracting the underlying firing rates from the spike trains of the RNN dataset. Shown are single trial examples for **d** GPFA, **e** vLGP, **f** PfLDS, and **g** LFADS. As can be seen by eye, the LFADS results are closer to the actual underlying rates than for the other models (black, firing rates of chaotic data RNN, red, inferred rates). **h-j** Summary *R*^2^ values between actual and inferred rates. Comparison using held-out data of the *R*^2^ values for **h** GPFA vs. LFADS, **i** vLGP vs. LFADS, and **j** PfLDS vs. LFADS. In all comparisons, LFADS yields a better fit to the data, for every single neuron.

**Figure 4:**
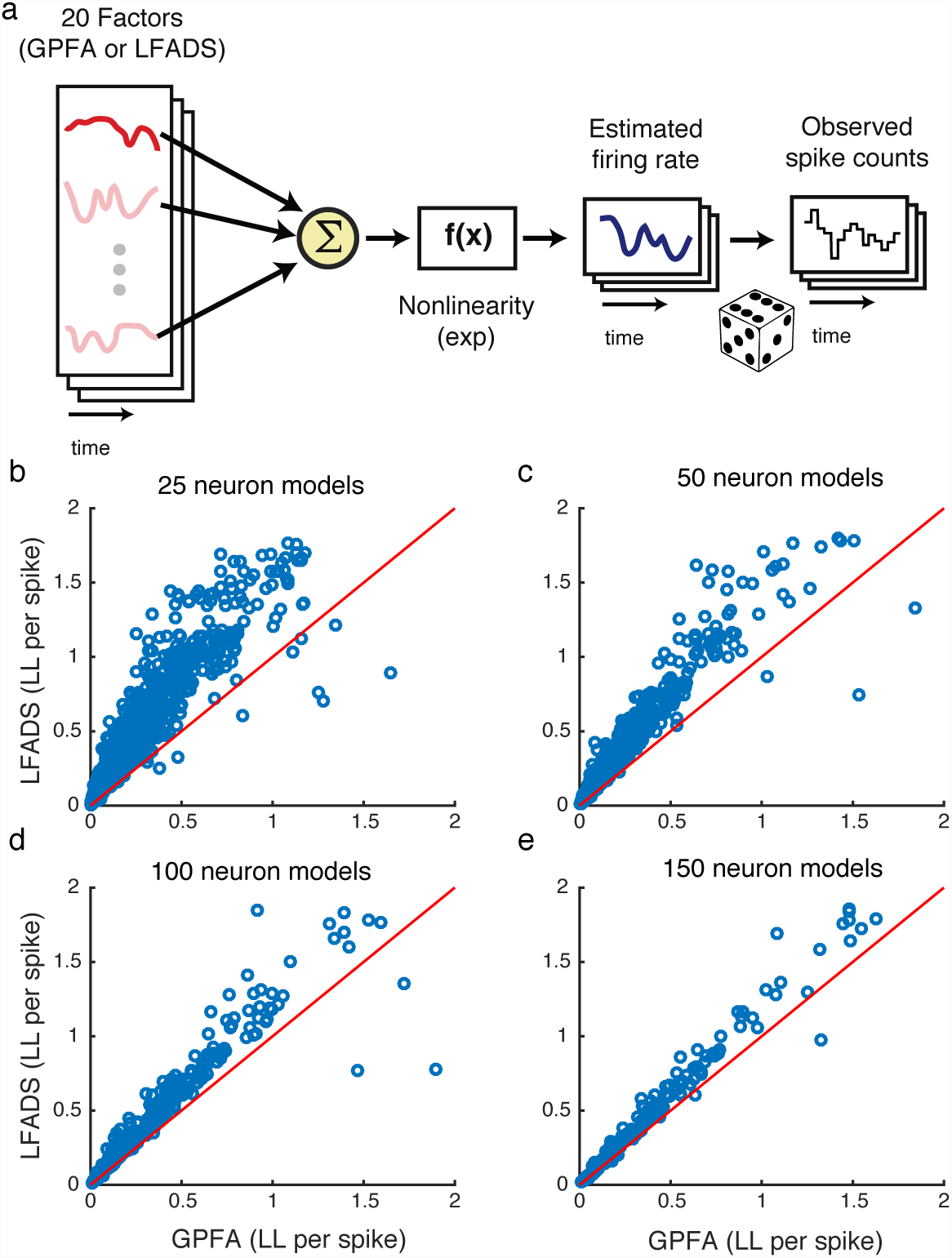
LFADS outperforms GPFA in predicting held-out neurons. When estimating latent state from real neural data, it is difficult to compare the accuracy of the estimates produced by different methods as the true neural latent state is unknown. However, if the latent state estimates produced by a technique are informative, then they may accurately describe held-out neural data, i.e., neurons that were simultaneously recorded but not used to train the methods. We compared the accuracy of LFADS against GPFA in predicting held-out neurons in the Maze dataset (**Figs. 2**, main text). As shown in **Fig. 2**, we sub-sampled neurons from the complete neural population (202 neurons total), and used the sub-sampled populations to estimate latent dynamics (e.g. 25, 50, 100, or 150 neurons to fit either LFADS or GPFA latent models). **a** We used a standard Generalized Linear Model (GLM) framework [4] to map the latent state estimates produced by LFADS or GPFA onto the binned spike counts (20 ms bins) for the remaining held-out neurons, e.g., for a model trained with 25 neurons, there are 177=202-25 held out neurons. **b-e** We then measured the improvement produced by the LFADS latent estimates over GPFA (evaluated using log likelihood per spike, LLPS [5]). For a given held-out neuron, we predicted the neuron’s firing rate based on the GLM fit, for all trials that were held out from the GLM fit. We then evaluated the LLPS of the observed spike trains given the predicted firing rates. We did this for held out neuron populations of **b** 25, **c** 50, **d** 100, or **e** 150 neurons. For almost all held-out neurons, LFADS-inferred latent state estimates were much more predictive about the spike counts of the held-out neurons than estimates produced by GPFA.

**Figure 5:**
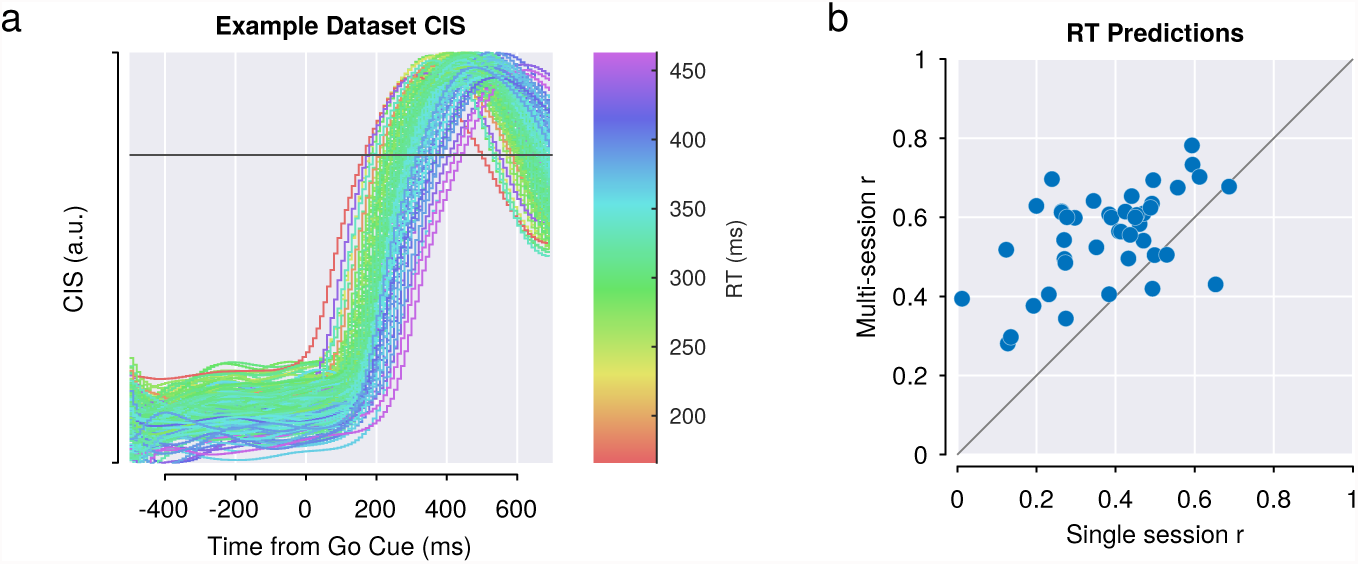
Dynamically stitched multi-session LFADS model outperforms single-session models in predicting reaction times. As defined in [2], the condition-independent signal (CIS) is a high variance component of motor cortical population activity obtained via demixing principal components analysis (DPCA [3]). The authors of [2] also demonstrated that threshold crossing time of the CIS on single trials is an effective predictor of reach reaction time (RT). Here we identify the CIS as a linear projection of LFADS factor trajectories. We apply DPCA to the factor outputs of each single-session and the multi-session LFADS models to identify the largest condition-independent component, and then threshold the CIS to predict RT on single trials. (a) Plot of condition-independent signals (CIS) for an example dataset. Each trace represents the CIS timecourse on a single trial, and is colored by that trial’s actual RT. **(b)** Plot of correlations between CIS-predicted RT and actual RT for multi-session LFADS vs. single-session LFADS models. Each point represents an individual recording session. A single CIS projection was computed for the multi-session model and applied for all sessions, whereas individual CIS projections were obtained for each single-session model.

**Figure 6:**
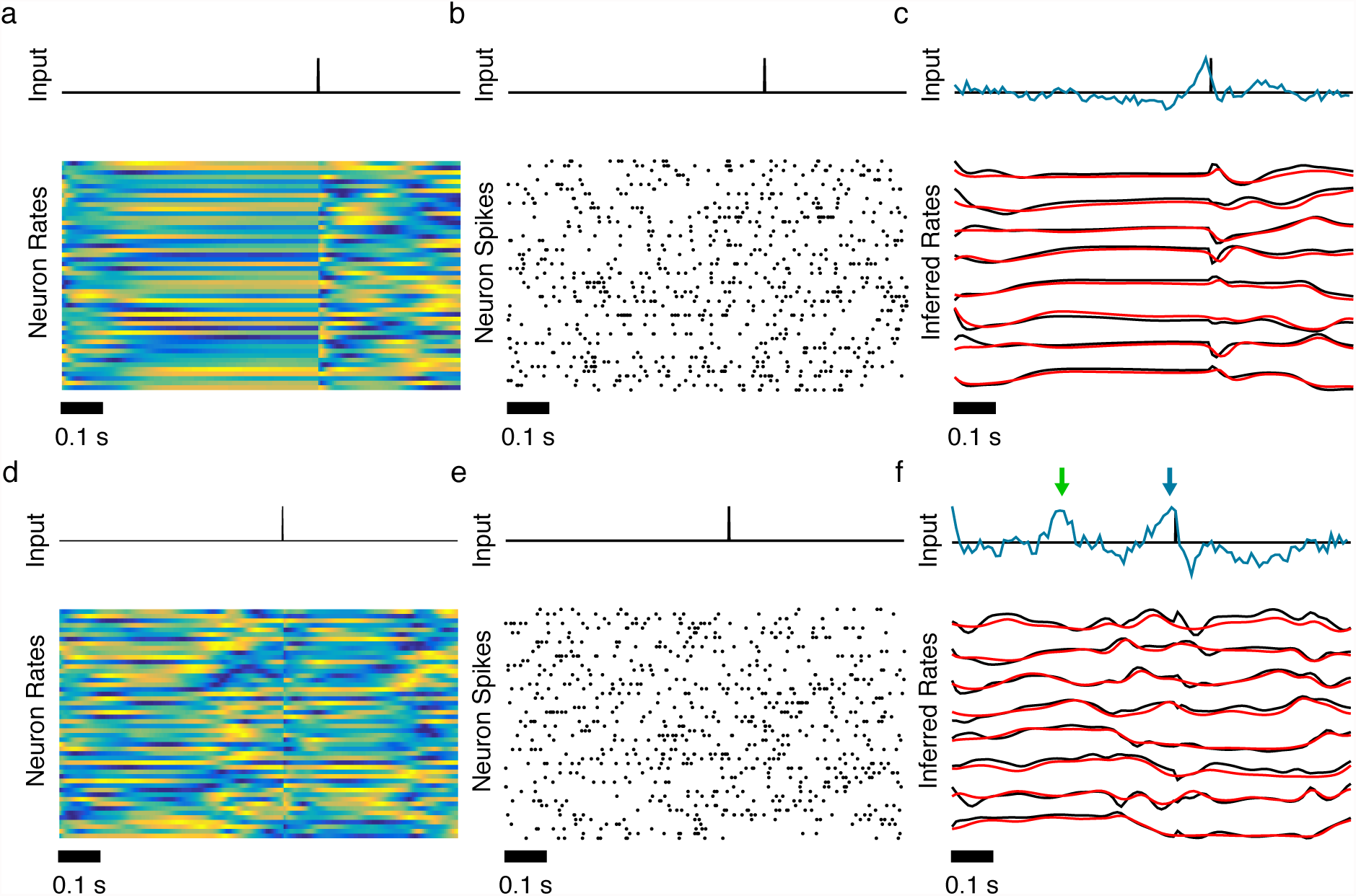
Inferring inputs from a chaotic data RNN with delta pulse inputs. We tested the ability of LFADS to infer the input to a dynamical system, specifically chaotic data RNNs, as used in the previous figure. During each trial, we perturbed the network by delivering a delta pulse at a random time *t*_pulse_ between 0.25s and 0.75s. The full trial length was 1s. This pulse affected the underlying rates produced by the data RNN, which subsequently affected the generated spike trains that were used as input to LFADS. To test the ability of the LFADS model to infer the timing of these input pulses, we included in the LFADS model an inferred input with dimensionality of 1. We explored two levels of dynamical complexity in the data RNNs (see Online Methods), defined by two values, 1.5 and 2.5, of a hyper-parameter to the data RNN, *γ*. **a-c** *γ* = 1.5. This value of *γ* value produces “gentler” chaotic activity in the data RNN than the higher value. **a** Example trial illustrating results from the *γ* = 1.5 chaotic data RNN with an external input (shown in black at the top of each column). Firing rates for the 50 simulated neurons. **b** Poisson-generated spike times for the simulated neurons. **c** Example trial showing (top) the actual (black) and inferred (cyan) input, and (bottom) actual firing rates of a subset of neurons in black and the corresponding inferred firing rates in red (bottom). **d-f** Same as a-c, but for *γ* = 2.5, which produces significantly more chaotic dynamics than *γ* = 1.5. **g** For this more difficult case, LFADS inferred the correct input (blue arrow), but also used the input to shape the dynamics at times there was no actual input (green arrow).

**Figure 7:**
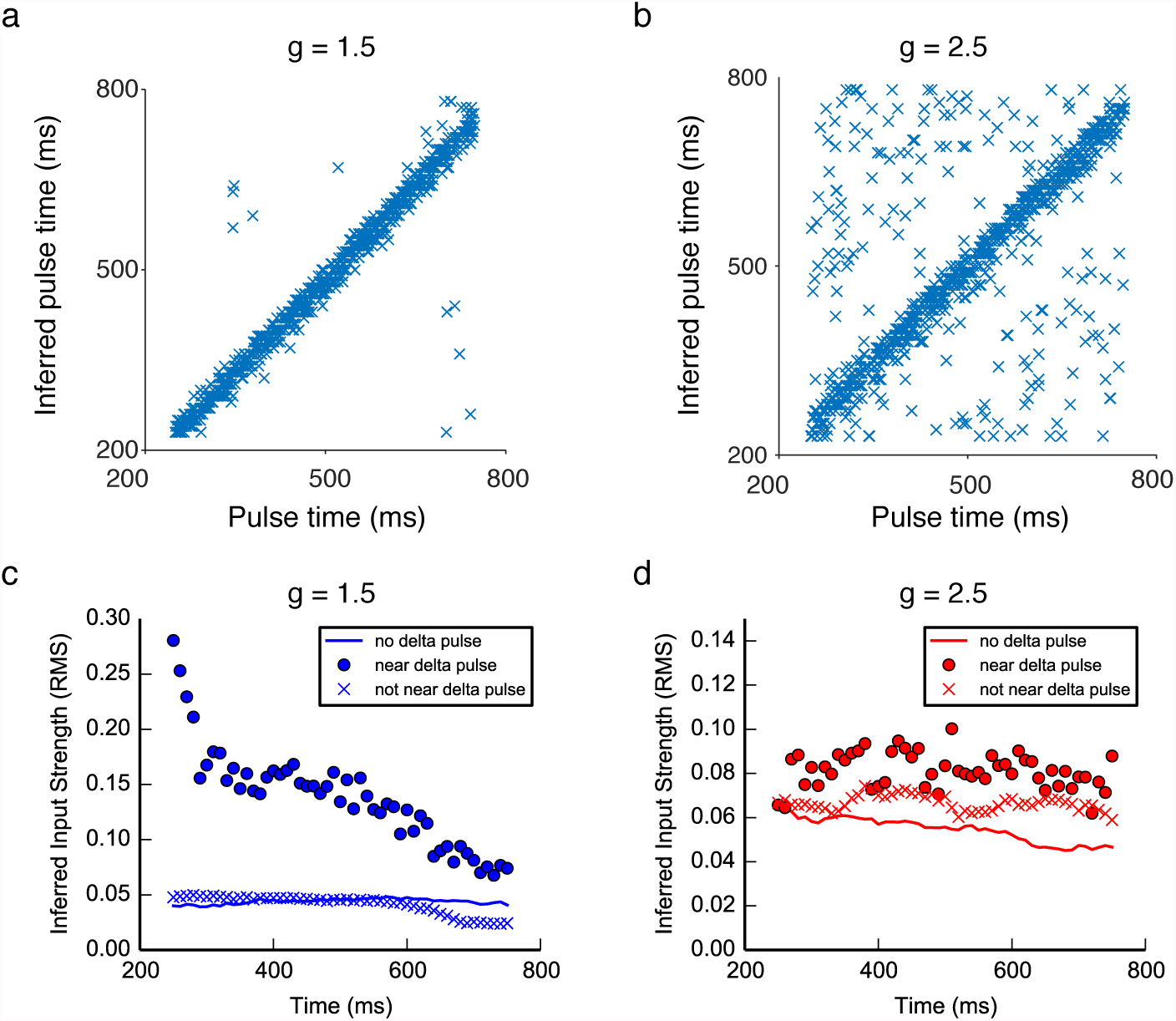
Summary of results of chaotic data RNNs receiving input pulses. We extracted averaged inferred inputs, 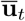, from the LFADS model (see Online Methods). **a,b** To see how this was related to the actual input pulse, we determined the time at which 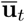 reached its maximum value. The time of the inferred input (time of the maximum of 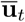; vertical axis) plotted against the actual time of the delta pulse (horizontal axis) for all trials. **a** Results using *γ* = 1.5, which induces chaotic dynamics. **b** and results using *γ* = 2.5, which induce strongly chaotic dynamics. These plots show that for the majority of trials, despite complex internal dynamics, LFADS was able to infer the correct timing of a strong input. However, LFADS did a better job of inferring the inputs in the case of simpler dynamics for two reasons. First, in the case of *γ* = 2.5, the complex dynamics reduces the effective magnitude of the perturbation caused by the input. Second, LFADS used the inferred input more actively to account for the non-input-driven dynamics as well as the input driven dynamics. We include this example of a highly chaotic data RNN to highlight the subtlety of interpreting an inferred input. **c-d** One possibility in using LFADS with inferred inputs (i.e. dimensionality of **u***_t_* ≥ 1) is that the data to be modeled is actually generated by an autonomous system, yet one, not knowing this fact, allows for an inferred input in LFADS. To study this case we utilized the four chaotic data RNNs described above, i.e. *γ* = 1.5, and *γ* = 2.5, with and without delta pulse inputs. We trained an LFADS model for each of the four cases, with an inferred input of dimensionality 1, despite the fact that two of the four data RNNs generated their data autonomously. After training we examined the strength of the average inferred input, 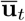, for each LFADS model. Our definition of strength is root-mean-square of the inferred input, averaged over an appropriate time window, 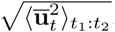. The results are show in panel **c** for the *γ* = 1.5 networks and panel **d** for the *γ* = 2.5 networks. The solid lines show the strength 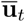 at each time point, for the data RNN that received no delta pulses, averaged across all examples. The ‘o’ and ‘x’ show the strength of 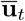 for the data RNN that received delta pulses, averaged in a time window around *t*, and averaged over all examples. Intuitively, a ‘o’ is the strength of 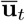 around a delta pulse at time *t*, and an ‘x’ is the strength of 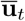 if there was no delta pulse around time *t*. Importantly, the strength of the inferred input when pulses were not present in the data was similar to the magnitude of inferred input when pulses were present in the data but not in the specific window. Further, when inputs were present in the data and within the specific window, the magnitude of the inferred input was higher on average than cases without inputs.

**Figure 8:**
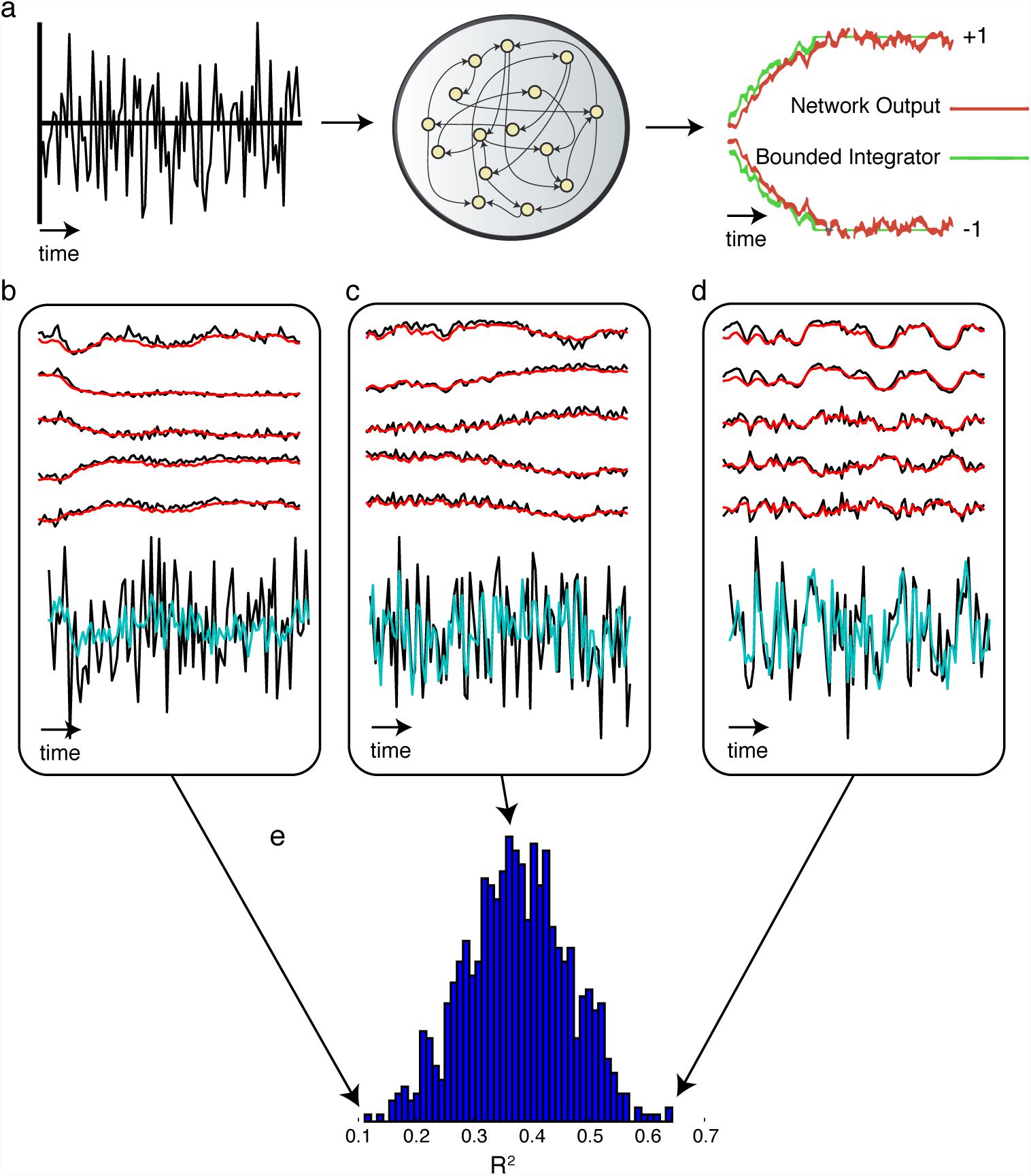
Inferring input from an integrate-to-bound data RNN. LFADS is able to model simulated neurons integrating a noisy input, and infer the noise signal itself. **a** Overview of the integration to bound task. On each trial, the data RNN receives noise drawn from a Gaussian distribution with mean 0, variance 0.0625. We trained an RNN to integrate this stochastic, 1-dimensional input to either a high (+1) or low (−1) bound. After the data RNN learned the task, we generated spiking data from 50 neurons using similar methodology as **Supp. Fig. 3** and fit an LFADS model to this data. **b-d** We fit an LFADS model to the data using 3200 1-second training examples, and evaluated its performance on 800 held-out trials. LFADS was able to accurately infer the ground truth firing rates (LFADS in red, ground truth in black). LFADS also inferred the associated white-noise input to the data RNN (LFADS cyan, ground truth in black, posterior means averaged over 1024 samples). These panels show the trials with the worst, median, and best measured *R*^2^ values between true and inferred inputs. **b** Trial with worst *R*^2^ = 0.11, **c** median *R*^2^ = 0.38, **d** and the best *R*^2^ = 0.64. **e** Histogram showing distribution of *R*^2^ values between true and inferred inputs for the 800 held-out trials.

**Figure 9:**
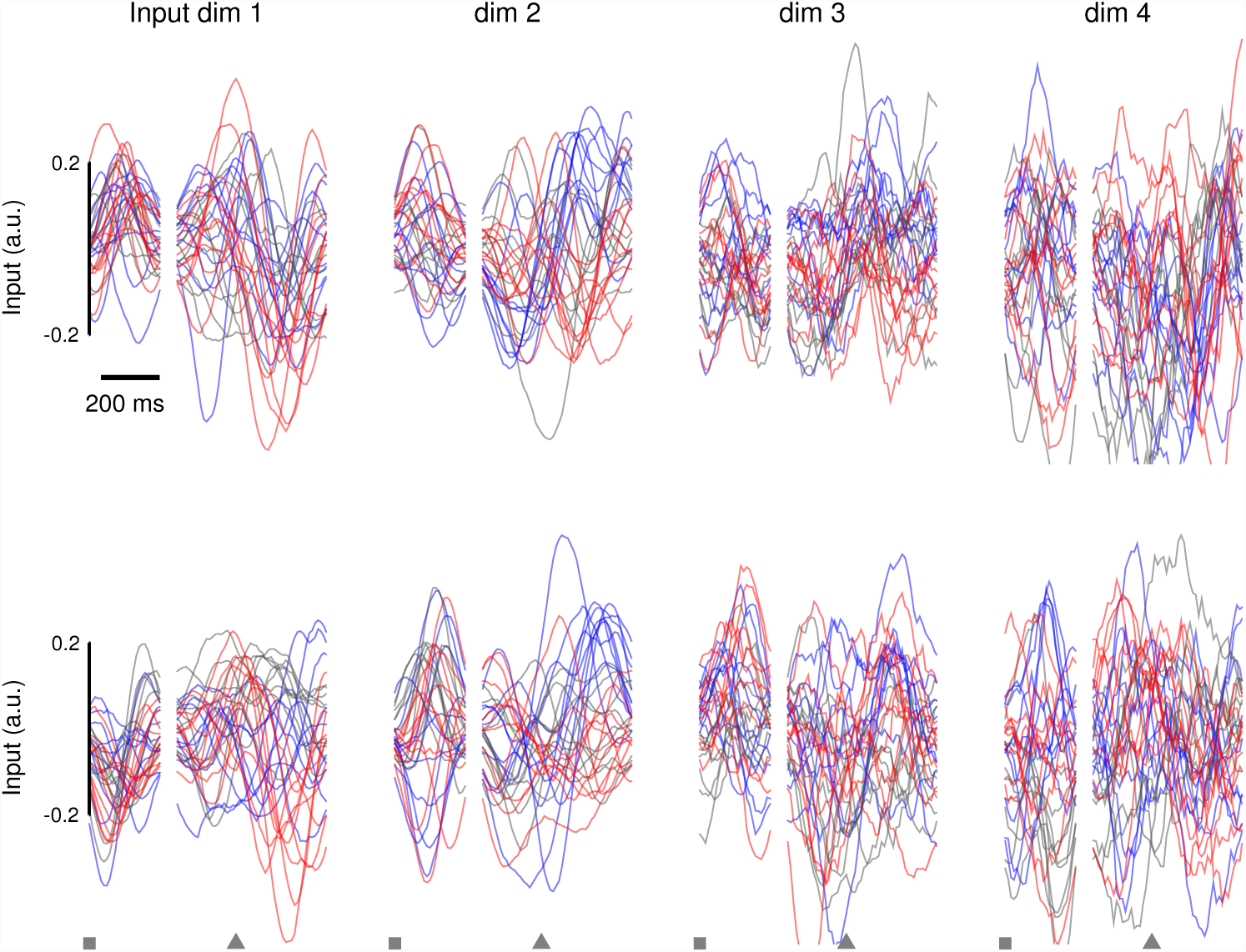
Inferred inputs from individual trials in the CursorJump task. This figure parallels **Fig. 6c** from the main text, with inferred inputs plotted for individual trials. To increase visibility, only 10 trials for each condition (reach direction and perturbation type) are shown. Individual traces were smoothed with an acausal Gaussian filter (60 ms s.d.). Despite the high variance across individual trials, several of the trends in the inferred inputs described in Fig 6 in the main text are visible at the single trial level. The inputs show information about the target of the upcoming reach on a single trial level, though individual traces are noisy. Specifically, for example, at the time of target onset (squares), the inferred input dimension 1 diverges for up vs. down reaches, but not for different perturbation types (as the information about perturbation type is not yet known at that phase of the task). Further, the inputs show information about the perturbation timing and identity on a single trial level, though again, individual traces are noisy. Specifically, around the time of the perturbation (arrow), the traces diverge for left-perturbed vs. right-perturbed vs. unperturbed trials (e.g., seen in dimension 2). Though these individual traces are noisy, Fig. 6d in the main text shows that these inputs can largely be separated on a single trial basis using a nonlinear dimensionality reduction algorithm, t-SNE.

**Video 1: LFADS reveals consistent rotational dynamics on individual trials.** Video contains two sequential movies showing the trajectories in neural population state space during individual reach trials for Monkey J (Fig 3 in the main text). The first movie illustrates the single-trial trajectories uncovered by smoothing the data with a Gaussian kernel. The second movie illustrates single-trial trajectories uncovered by LFADS. 2296 trials are shown, representing 108 conditions of the “maze” dataset.

**Video 2: Mult-session LFADS finds consistent representations for individual trials across sessions.** Video contains four sequential movies showing the trajectories in neural population state space during individual reach trials for Monkey P (Fig 5 in the main text). The first movie illustrates the single-trial trajectories for a single session uncovered by smoothing the data and using the principal components regression technique across all sessions (Online Methods). Each individual trial is aligned to the time that movement was detected (movement onset), and the movie covers the timeframe from 350 ms before to 225 ms after movement onset. Colors represent 7 reach directions. The second movie illustrates single-trial trajectories for all trials over 42 / 44 sessions (2 were omitted for ease of presentation). Each panel denotes a different recording session. The third movie illustrates single-trial trajectories uncovered by multi-session LFADS for the first session, and the fourth movie illustrates single-trial trajectorues for 42 sessions. Multi-session movies include approximately 14,500 trials, 38 separate electrode penetration sites and spanned 162 days from the first to the last session.

## References

Afshar, Afsheen, Gopal Santhanam, Byron M. Yu, Stephen I. Ryu, Maneesh Sahani, and Krishna V. Shenoy. 2011. “Single-Trial Neural Correlates of Arm Movement Preparation.” Neuron 71 (3): 555–64.

Aghagolzadeh, Mehdi, and Wilson Truccolo. 2014. “Latent State-Space Models for Neural Decoding.” Conference Proceedings:… Annual International Conference of the IEEE Engineering in Medicine and Biology Society. IEEE Engineering in Medicine and Biology Society. Conference 2014: 3033–36.

Ahrens, Misha B., Jennifer M. Li, Michael B. Orger, Drew N. Robson, Alexander F. Schier, Florian Engert, and Ruben Portugues. 2012. “Brain-Wide Neuronal Dynamics during Motor Adaptation in Zebrafish.” Nature 485 (7399): 471–77.

Carnevale, Federico, Victor de Lafuente, Ranulfo Romo, Omri Barak, and Néstor Parga. 2015. “Dynamic Control of Response Criterion in Premotor Cortex during Perceptual Detection under Temporal Uncertainty.” Neuron 86 (4): 1067–77.

Churchland, Mark M., John P. Cunningham, Matthew T. Kaufman, Justin D. Foster, Paul Nuyujukian, Stephen I. Ryu, and Krishna V. Shenoy. 2012. “Neural Population Dynamics during Reaching.” Nature 487 (7405): 51–56.

Donoghue, J. P., J. N. Sanes, N. G. Hatsopoulos, and G. Gaál. 1998. “Neural Discharge and Local Field Potential Oscillations in Primate Motor Cortex during Voluntary Movements.” Journal of Neurophysiology 79 (1): 159–73.

Gao, Peiran, and Surya Ganguli. 2015. “On Simplicity and Complexity in the Brave New World of Large-Scale Neuroscience.” Current Opinion in Neurobiology 32 (June). Elsevier: 148–55.

Gao, Yuanjun, Evan W. Archer, Liam Paninski, and John P. Cunningham. 2016. “Linear Dynamical Neural Population Models through Nonlinear Embeddings.” In Advances in Neural Information Processing Systems 29, edited by D. D. Lee, M. Sugiyama, U. V. Luxburg, I. Guyon, and R. Garnett, 163–71. Curran Associates, Inc.

Gilja, Vikash, Chethan Pandarinath, Christine H. Blabe, Paul Nuyujukian, John D. Simeral, Anish A. Sarma, Brittany L. Sorice, et al. 2015. “Clinical Translation of a High-Performance Neural Prosthesis.” Nature Medicine 21 (10): 1142–45.

Gregor, Karol, Ivo Danihelka, Alex Graves, Danilo Jimenez Rezende, and Daan Wierstra. 2015. “DRAW: A Recurrent Neural Network For Image Generation.” arXiv [cs.CV]. arXiv. http://arxiv.org/abs/1502.04623.

Harvey, Christopher D., Philip Coen, and David W. Tank. 2012. “Choice-Specific Sequences in Parietal Cortex during a Virtual-Navigation Decision Task.” Nature 484 (7392): 62–68.

Kao, Jonathan C., Paul Nuyujukian, Stephen I. Ryu, Mark M. Churchland, John P. Cunningham, and Krishna V. Shenoy. 2015. “Single-Trial Dynamics of Motor Cortex and Their Applications to Brain-Machine Interfaces.” Nature Communications 6 (July): 7759.

Kato, Saul, Harris S. Kaplan, Tina Schrödel, Susanne Skora, Theodore H. Lindsay, Eviatar Yemini, Shawn Lockery, and Manuel Zimmer. 2015. “Global Brain Dynamics Embed the Motor Command Sequence of Caenorhabditis Elegans.” Cell 163 (3). Elsevier: 656–69.

Kaufman, Matthew T., Mark M. Churchland, Stephen I. Ryu, and Krishna V. Shenoy. 2014. “Cortical Activity in the Null Space: Permitting Preparation without Movement.” Nature Neuroscience 17 (3): 440–48.

Kaufman, Matthew T., Jeffrey S. Seely, David Sussillo, Stephen I. Ryu, Krishna V. Shenoy, and Mark M. Churchland. 2016. “The Largest Response Component in the Motor Cortex Reflects Movement Timing but Not Movement Type.” eNeuro 3 (4). eneuro.org. doi:10.1523/ENEURO.0085-16.2016.

Kingma, Diederik P., and Max Welling. 2013. “Auto-Encoding Variational Bayes.” arXiv [stat.ML]. arXiv. http://arxiv.org/abs/1312.6114v10.

Kobak, Dmitry, Wieland Brendel, Christos Constantinidis, Claudia E. Feierstein, Adam Kepecs, Zachary F. Mainen, Xue-Lian Qi, Ranulfo Romo, Naoshige Uchida, and Christian K. Machens. 2016a. “Demixed Principal Component Analysis of Neural Population Data.” eLife 5 (April). doi:10.7554/eLife.10989.

Linderman, Scott, Matthew Johnson, Andrew Miller, Ryan Adams, David Blei, and Liam Paninski. 2017. “Bayesian Learning and Inference in Recurrent Switching Linear Dynamical Systems.” In Artificial Intelligence and Statistics, 914–22. proceedings.mlr.press.

Macke, Jakob H., Lars Buesing, John P. Cunningham, Byron M. Yu, Krishna V. Shenoy, and Maneesh Sahani. 2011. “Empirical Models of Spiking in Neural Populations.” In Advances in Neural Information Processing Systems, 1350–58.

Mante, Valerio, David Sussillo, Krishna V. Shenoy, and William T. Newsome. 2013. “Context-Dependent Computation by Recurrent Dynamics in Prefrontal Cortex.” Nature 503 (7474): 78–84.

Murthy, V. N., and E. E. Fetz. 1996. “Synchronization of Neurons during Local Field Potential Oscillations in Sensorimotor Cortex of Awake Monkeys.” Journal of Neurophysiology 76 (6): 3968–82.

Pandarinath, Chethan, Vikash Gilja, Christine H. Blabe, Paul Nuyujukian, Anish A. Sarma, Brittany L. Sorice, Emad N. Eskandar, Leigh R. Hochberg, Jaimie M. Henderson, and Krishna V. Shenoy. 2015. “Neural Population Dynamics in Human Motor Cortex during Movements in People with ALS.” eLife 4 (June): e07436.

Pandarinath, Chethan, Paul Nuyujukian, Christine H. Blabe, Brittany L. Sorice, Jad Saab, Francis R. Willett, Leigh R. Hochberg, Krishna V. Shenoy, and Jaimie M. Henderson. 2017. “High Performance Communication by People with Paralysis Using an Intracortical Brain-Computer Interface.” eLife 6 (February). doi:10.7554/eLife.18554.

Petreska, Biljana, Byron M. Yu, John P. Cunningham, Gopal Santhanam, Stephen I. Ryu, Krishna V. Shenoy, and Maneesh Sahani. 2011. “Dynamical Segmentation of Single Trials from Population Neural Data.” In Advances in Neural Information Processing Systems 24, edited by J. Shawe-Taylor, R. S. Zemel, P. L. Bartlett, F. Pereira, and K. Q. Weinberger, 756–64. Curran Associates, Inc.

Sadtler, Patrick T., Kristin M. Quick, Matthew D. Golub, Steven M. Chase, Stephen I. Ryu, Elizabeth C. Tyler-Kabara, Byron M. Yu, and Aaron P. Batista. 2014. “Neural Constraints on Learning.” Nature 512 (7515): 423–26.

Salinas, E., and L. F. Abbott. 1994. “Vector Reconstruction from Firing Rates.” Journal of Computational Neuroscience 1 (1-2): 89–107.

Sussillo, David, Paul Nuyujukian, Joline M. Fan, Jonathan C. Kao, Sergey D. Stavisky, Stephen Ryu, and Krishna Shenoy. 2012. “A Recurrent Neural Network for Closed-Loop Intracortical Brain-Machine Interface Decoders.” Journal of Neural Engineering 9 (2): 026027.

Sussillo, David, Sergey D. Stavisky, Jonathan C. Kao, Stephen I. Ryu, and Krishna V. Shenoy. 2016. “Making Brain-machine Interfaces Robust to Future Neural Variability.” Nature Communications 7: 13749.

Willett, Francis R., Chethan Pandarinath, Beata Jarosiewicz, Brian A. Murphy, William D. Memberg, Christine H. Blabe, Jad Saab, et al. 2017. “Feedback Control Policies Employed by People Using Intracortical Brain-Computer Interfaces.” Journal of Neural Engineering 14 (1): 016001.

Yu, Byron M., John P. Cunningham, Gopal Santhanam, Stephen I. Ryu, Krishna V. Shenoy, and Maneesh Sahani. 2009. “Gaussian-Process Factor Analysis for Low-Dimensional Single-Trial Analysis of Neural Population Activity.” Journal of Neurophysiology 102 (1): 614–35.

Yuste, Rafael. 2015. “From the Neuron Doctrine to Neural Networks.” Nature Reviews. Neuroscience 16 (8): 487–97.

Zhao, Yuan, and Il Memming Park. 2017. “Variational Latent Gaussian Process for Recovering Single-Trial Dynamics from Population Spike Trains.” Neural Computation 29 (5): 1293–1316.

## References

[1] Bayer, J., and Osendorfer, C. Learning stochastic recurrent networks. arXiv preprint arXiv:1411.7610 (2014).

[2] Bowman, S. R., Vilnis, L., Vinyals, O., Dai, A. M., Jozefowicz, R., and Bengio, S. Generating sentences from a continuous space. Conference on Computational Natural Language Learning (CoNLL) (2016).

[3] Carnevale, F., de Lafuente, V., Romo, R., Barak, O., and Parga, N. Dynamic control of response criterion in premotor cortex during perceptual detection under temporal uncertainty. Neuron 86, 4 (2015), 1067–1077.

[4] Chung, J., Gulcehre, C., Cho, K., and Bengio, Y. Empirical evaluation of gated recurrent neural networks on sequence modeling. arXiv preprint arXiv:1412.3555 (2014).

[5] Chung, J., Kastner, K., Dinh, L., Goel, K., Courville, A., and Bengio, Y. A recurrent latent variable model for sequential data. In Advances in Neural Information Processing Systems (NIPS) (2015).

[6] Churchland, M. M., Cunningham, J. P., Kaufman, M. T., Foster, J. D., Nuyujukian, P., Ryu, S. I., and Shenoy, K. V. Neural population dynamics during reaching. Nature 487, 7405 (2012), 51–56.

[7] Fan, J. M., Nuyujukian, P., Kao, J. C., Chestek, C. A., Ryu, S. I., and Shenoy, K. V. Intention estimation in brain–machine interfaces. Journal of neural engineering 11, 1 (2014), 016004.

[8] Gao, P., and Ganguli, S. On simplicity and complexity in the brave new world of large-scale neuroscience. Current opinion in neurobiology 32 (2015), 148–155.

[9] Gao, Y., Archer, E., Paninski, L., and Cunningham, J. P. Linear dynamical neural population models through nonlinear embeddings. arXiv preprint arXiv:1605.08454 (2016).

[10] Gao, Y., Busing, L., Shenoy, K. V., and Cunningham, J. P. High-dimensional neural spike train analysis with generalized count linear dynamical systems. In Advances in Neural Information Processing Systems (2015), pp. 2044–2052.

[11] Gilja, V., Nuyujukian, P., Chestek, C. A., Cunningham, J. P., Byron, M. Y., Fan, J. M., Churchland, M. M., Kaufman, M. T., Kao, J. C., Ryu, S. I., et al. A high-performance neural prosthesis enabled by control algorithm design. Nature neuroscience 15, 12 (2012), 1752–1757.

[12] Gilja, V., Pandarinath, C., Blabe, C. H., Nuyujukian, P., Simeral, J. D., Sarma, A. A., Sorice, B. L., Perge, J. A., Jarosiewicz, B., Hochberg, L. R., et al. Clinical translation of a high-performance neural prosthesis. Nature medicine 21, 10 (2015), 1142–1145.

[13] Gregor, K., Danihelka, I., Graves, A., Rezende, D. J., and Wierstra, D. Draw: A recurrent neural network for image generation. arXiv preprint arXiv:1502.04623 (2015).

[14] Hinton, G. E., Srivastava, N., Krizhevsky, A., Sutskever, I., and Salakhutdinov, R. R. Improving neural networks by preventing co-adaptation of feature detectors. arXiv preprint arXiv:1207.0580 (2012).

[15] Karl, M., Soelch, M., Bayer, J., and van der Smagt, P. Deep variational bayes filters: Unsupervised learning of state space models from raw data. arXiv preprint arXiv:1605.06432 (2016).

[16] Kato, S., Kaplan, H. S., Schrödel, T., Skora, S., Lindsay, T. H., Yemini, E., Lockery, S., and Zimmer, M. Global brain dynamics embed the motor command sequence of caenorhabditis elegans. Cell 163, 3 (2015), 656–669.

[17] Kaufman, M. T., Seely, J. S., Sussillo, D., Ryu, S. I., Shenoy, K. V., and Churchland, M. M. The largest response component in the motor cortex reflects movement timing but not movement type. eneuro 3, 4 (2016), ENEURO–0085.

[18] Kaufman, M. T., Seely, J. S., Sussillo, D., Ryu, S. I., Shenoy, K. V., and Churchland, M. M. The largest response component in the motor cortex reflects movement timing but not movement type. eNeuro 3, 4 (July 2016).

[19] Kingma, D. P., and Welling, M. Auto-encoding variational bayes. In Proceedings of the 2nd International Conference on Learning Representations (ICLR) (2013), no. 2014.

[20] Kobak, D., Brendel, W., Constantinidis, C., Feierstein, C. E., Kepecs, A., Mainen, Z. F., Qi, X.-L., Romo, R., Uchida, N., and Machens, C. K. Demixed principal component analysis of neural population data. eLife 5 (12 Apr. 2016).

[21] Krishnan, R. G., Shalit, U., and Sontag, D. Deep kalman filters. arXiv preprint arXiv:1511.05121 (2015).

[22] Maaten, L. v. d., and Hinton, G. Visualizing data using t-sne. Journal of Machine Learning Research 9, Nov (2008), 2579–2605.

[23] Macke, J. H., Buesing, L., Cunningham, J. P., Yu, B. M., Shenoy, K. V., and Sahani, M. Empirical models of spiking in neural populations. In Advances in neural information processing systems (2011), pp. 1350–1358.

[24] Mante, V., Sussillo, D., Shenoy, K. V., and Newsome, W. T. Context-dependent computation by recurrent dynamics in prefrontal cortex. Nature 503, 7474 (2013), 78–84.

[25] Pandarinath, C., Gilja, V., Blabe, C. H., Nuyujukian, P., Sarma, A. A., Sorice, B. L., Eskandar, E. N., Hochberg, L. R., Henderson, J. M., and Shenoy, K. V. Neural population dynamics in human motor cortex during movements in people with als. Elife 4 (2015), e07436.

[26] Pandarinath, C., Nuyujukian, P., Blabe, C. H., Sorice, B. L., Saab, J., Willett, F. R., Hochberg, L. R., Shenoy, K. V., and Henderson, J. M. High performance communication by people with paralysis using an intracortical brain-computer interface. eLife 6 (2017), e18554.

[27] Petreska, B., Byron, M. Y., Cunningham, J. P., Santhanam, G., Ryu, S. I., Shenoy, K. V., and Sahani, M. Dynamical segmentation of single trials from population neural data. In Advances in neural information processing systems (2011), pp. 756–764.

[28] Rajan, K., Harvey, C. D., and Tank, D. W. Recurrent network models of sequence generation and memory. Neuron 90 (2016), 1–15.

[29] Rezende, D. J., Mohamed, S., and Wierstra, D. Stochastic backpropagation and approximate inference in deep generative models. In International Conference on Machine Learning, 2014 (2014).

[30] Salinas, E., and Abbott, L. Vector reconstruction from firing rates. Journal of computational neuroscience 1, 1 (1994), 89–107.

[31] Sussillo, D., and Abbott, L. F. Generating coherent patterns of activity from chaotic neural networks. Neuron 63, 4 (2009), 544–557.

[32] Sussillo, D., Churchland, M. M., Kaufman, M. T., and Shenoy, K. V. A neural network that finds a naturalistic solution for the production of muscle activity. Nature neuroscience 18, 7 (2015), 1025–1033.

[33] Watter, M., Springenberg, J., Boedecker, J., and Riedmiller, M. Embed to control: A locally linear latent dynamics model for control from raw images. In Advances in Neural Information Processing Systems (2015), pp. 2746–2754.

[34] Willett, F. R., Pandarinath, C., Jarosiewicz, B., Murphy, B. A., Memberg, W. D., Blabe, C. H., Saab, J., Walter, B. L., Sweet, J. A., Miller, J. P., et al. Feedback control policies employed by people using intracortical brain–computer interfaces. Journal of Neural Engineering 14, 1 (2016), 016001.

[35] Yu, B. M., Cunningham, J. P., Santhanam, G., Ryu, S. I., Shenoy, K. V., and Sahani, M. Gaussian-process factor analysis for low-dimensional single-trial analysis of neural population activity. In Advances in neural information processing systems (2009), pp. 1881–1888.

[36] Zaremba, W., Sutskever, I., and Vinyals, O. Recurrent neural network regularization. arXiv preprint arXiv:1409.2329 (2014).

[37] Zhao, Y., and Park, I. M. Variational latent gaussian process for recovering single-trial dynamics from population spike trains. arXiv preprint arXiv:1604.03053 (2016).

## References

[1] Gao, Y., Archer, E., Paninski, L., and Cunningham, J. P. Linear dynamical neural population models through nonlinear embeddings. arXiv preprint arXiv:1605.08454 (2016).

[2] Kaufman, M. T., Seely, J. S., Sussillo, D., Ryu, S. I., Shenoy, K. V., and Churchland, M. M. The largest response component in the motor cortex reflects movement timing but not movement type. eneuro 3, 4 (2016), ENEURO–0085.

[3] Kobak, D., Brendel, W., Constantinidis, C., Feierstein, C. E., Kepecs, A., Mainen, Z. F., Qi, X.-L., Romo, R., Uchida, N., and Machens, C. K. Demixed principal component analysis of neural population data. eLife 5 (2016), e10989.

[4] Paninski, L. Maximum likelihood estimation of cascade point-process neural encoding models. Network: Computation in Neural Systems 15, 4 (2004), 243–262.

[5] Williamson, R. S., Sahani, M., and Pillow, J. W. The equivalence of information-theoretic and likelihood-based methods for neural dimensionality reduction. PLoS Comput Biol 11, 4 (2015), e1004141.

[6] Yu, B. M., Cunningham, J. P., Santhanam, G., Ryu, S. I., Shenoy, K. V., and Sahani, M. Gaussian-process factor analysis for low-dimensional single-trial analysis of neural population activity. In Advances in neural information processing systems (2009), pp. 1881–1888.

[7] Zhao, Y., and Park, I. M. Variational latent gaussian process for recovering single-trial dynamics from population spike trains. arXiv preprint arXiv:1604.03053 (2016).

